# Cell-Specific Transposable Element Gene Expression Analysis Identifies Associations with Systemic Lupus Erythematosus Phenotypes

**DOI:** 10.1101/2023.11.27.567477

**Authors:** Zachary Cutts, Sarah Patterson, Lenka Maliskova, Kimberly E. Taylor, Chun Ye, Maria Dall’Era, Jinoos Yazdany, Lindsey Criswell, Gabriela K. Fragiadakis, Charles Langelier, John A. Capra, Marina Sirota, Cristina M Lanata

**Author notes:** these authors contributed equally to this work (Marina Sirota, Cristina M Lanata).

## Abstract

There is an established yet unexplained link between interferon (IFN) and systemic lupus erythematosus (SLE). The expression of sequences derived from transposable elements (TEs) may contribute to production of type I IFNs and generation of autoantibodies. We profiled cell-sorted RNA-seq data (CD4+ T cells, CD14+ monocytes, CD19+ B cells, and NK cells) from PBMCs of 120 SLE patients and quantified TE expression identifying 27,135 TEs. We tested for differential TE expression across 10 SLE phenotypes including autoantibody production and disease activity and discovered 731 differentially expressed (DE) TEs whose effects were mostly cell-specific and phenotype-specific. DE TEs were enriched for specific families and viral genes encoded in TE sequences. Increased expression of DE TEs was associated with genes involved in antiviral activity such as LY6E, ISG15, TRIM22 and pathways such as interferon signaling. These findings suggest that expression of TEs contributes to activation of SLE-related mechanisms in a cell-specific manner, which can impact disease diagnostics and therapeutics.

## Introduction

Systemic lupus erythematosus (SLE) is a heterogeneous chronic autoimmune disease characterized by antibodies against nucleic acids and associated proteins(1, 2). There is also varying prevalence of clinical manifestations among different racial and ethnic groups, and the risk of developing severe manifestations is increased in African Americans, Asian/Pacific Islanders, and Hispanic patients(3). Patients that self-report as Asian experience disproportionate burden of SLE compared to White patients, including greater incidence and prevalence of SLE, higher disease activity, worse organ damage, and heightened mortality(4). Transcriptomic studies have been undertaken to characterize the heterogeneity in SLE and have found expression of interferon (IFN)-regulated genes to be associated with disease. Genetic studies have shown that the heterogeneity in SLE may be due in part to the presence of pathogenic disease associated variants in non-coding regions(5–7).

Transposable element (TE) derived sequences make up approximately 50% of the human genome. TEs are mobile genetic elements capable of changing their location within genomes. The vast majority of TE sequences in the human genome are inactive and can no longer transpose, but they still contain sequences with the potential to encode proteins and functional gene regulatory elements. A small number of specific classes of TEs are intact and active in humans, and there is some evidence of differences across human populations(8). As a result, TEs have roles both in human health and diseases such as cancer and autoimmunity(9–11). Recent work has suggested that two subsets of TEs, human endogenous retroviruses (HERVs) and long interspersed nuclear elements (LINEs), may play a pathogenic role in SLE(12, 13). One hypothesis for how TEs could contribute to autoimmunity is that expression of TEs activates an immune response when TEs produce nucleic acids or proteins that resemble pathogen-associated molecular patterns (PAMPs) of ancient exogenous viruses(14, 15).

In previous work, over 100 locus-specific HERVs were shown to be differentially expressed in SLE patients and correlate with lupus clinical parameters such as presence of dsDNA, anti–RNP, and anti-SM antibodies(16). In addition, HERV-E clone 4-1 mRNA expression correlated with SLEDAI score in CD4+ T cells(17). Previous studies have identified elevated expression of HERVs in SLE, implicating HERV-E clone 4-1 in PBMCs and HRES1/p28 in B cells(18, 19). In addition, more recent work has characterized TEs in blood from SLE patients compared to matched controls, and found upregulation of TEs in SLE(11, 16, 20). These studies also investigated whether TEs contribute to the IFN signature observed in SLE patients, with one study finding a positive correlation between HERV expression and the IFN signature, while the other did not(11, 16).

Although many studies have explored TE expression between SLE patients and controls, none have defined locus specific TE expression in immune cell types relevant to SLE or characterized the relationship between TE expression and different lupus manifestations in a diverse patient cohort. In addition, prior work has not examined the cell specific association of differentially expressed (DE) TEs in SLE with gene expression and gene set enrichment analysis. The goal of this study was to characterize the role of TEs in the clinical heterogeneity of SLE across multiple cell types. Here we report on TE expression in 4 distinct cell types and 10 SLE subphenotypes, carry out family and viral gene enrichment analysis and correlate cell-specific TE expression with SLE heterogeneity as well as cell-specific transcriptomics in a diverse cohort of lupus patients with extensive phenotypic data.

## Results

The study consists of 120 SLE participants from the California Lupus Epidemiology Study (CLUES) (Figure 1A). The majority of participants were female, with an age distribution between20 and 82 (Table 1). In order to capture cell type specific elements across major immune populations, we leveraged cell-sorted RNA-seq data (CD4+ T cells, CD14+ monocytes, B cells, and NK cells) from PBMCs of patients that were previously published by Andreoletti et al(2). After standard quality control measures, we quantified expression of 27,135 HERVs and LINEs (TEs) in 4 cell-types using Telescope(10). We found that both TE and gene expression is cell specific in lupus patients as observed by the clear clustering in principal component analyses (PCA) of both TE expression (Figure 1B) and gene expression (Figure 1C).

**Figure 1.**
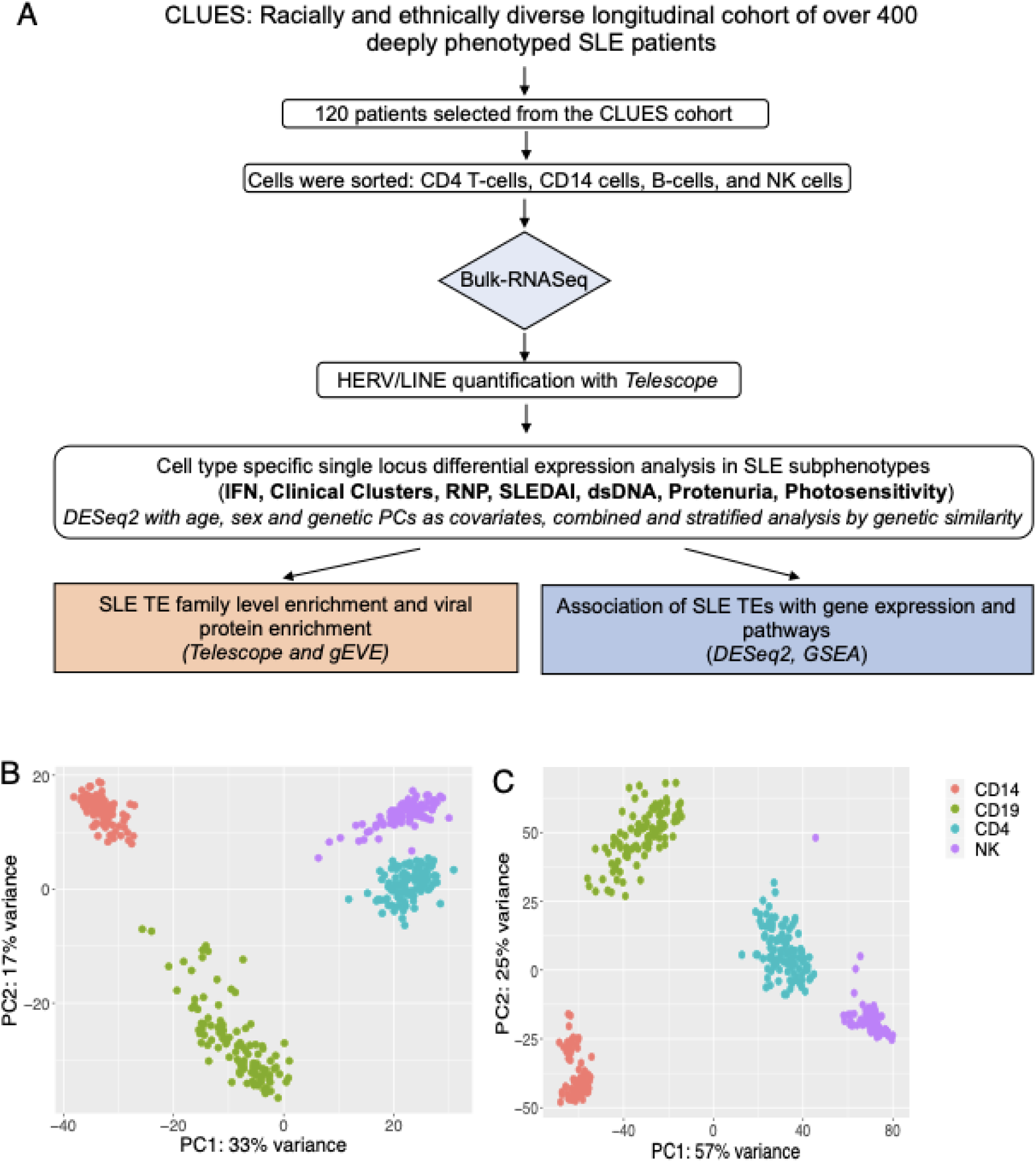
Study Overview and Comparison of Gene and TE expression between cell types. A. 120 patients selected for cell sorted bulk RNA-seq analyses from the CLUES cohort. Data were QC’ed and HERVs and LINE were quantified using Telescope. DESeq2 was used for cell specific differential expression of locus specific TEs for SLE subphenotypes. Cell specific differentially expressed TEs from all SLE subphenotypes were used to perform family enrichment, TE derived viral protein enrichment, and association with gene expression and pathway analysis. B. TE expression is cell specific in lupus - PCA plot based visualization colored by cell type. C. Gene expression PCA plot based visualization colored by cell type

**Table 1.**
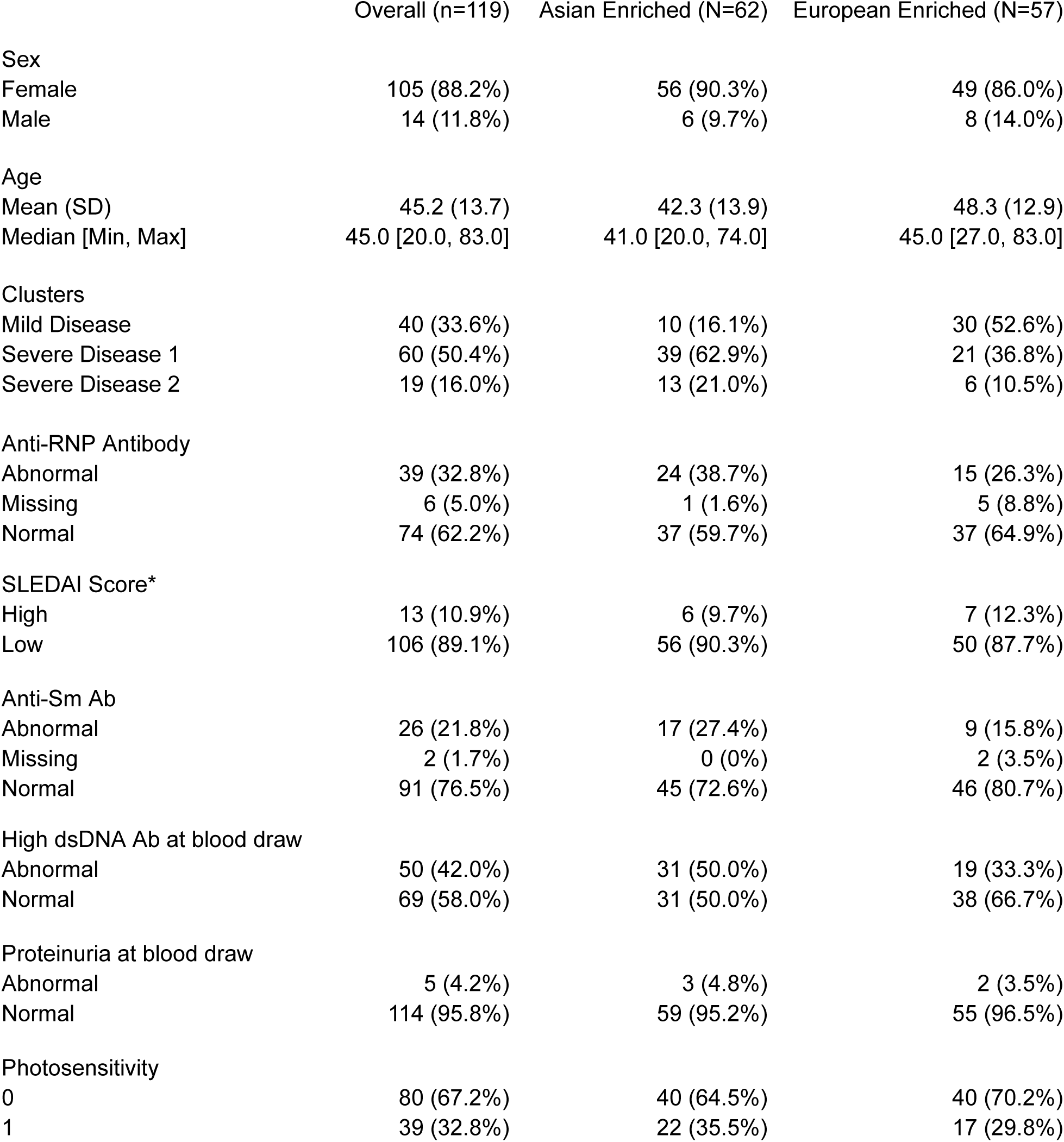
Cohort Characteristics.

|  | Overall (n=119) | Asian Enriched (N=62) | European Enriched (N=57) |
| --- | --- | --- | --- |
| Sex |  |  |  |
| Female | 105 (88.2%) | 56 (90.3%) | 49 (86.0%) |
| Male | 14 (11.8%) | 6 (9.7%) | 8 (14.0%) |
| Age |  |  |  |
| Mean (SD) | 45.2 (13.7) | 42.3 (13.9) | 48.3 (12.9) |
| Median [Min, Max] | 45.0 [20.0, 83.0] | 41.0 [20.0, 74.0] | 45.0 [27.0, 83.0] |
| Clusters |  |  |  |
| Mild Disease | 40 (33.6%) | 10 (16.1%) | 30 (52.6%) |
| Severe Disease 1 | 60 (50.4%) | 39 (62.9%) | 21 (36.8%) |
| Severe Disease 2 | 19 (16.0%) | 13 (21.0%) | 6 (10.5%) |
| Anti-RNP Antibody |  |  |  |
| Abnormal | 39 (32.8%) | 24 (38.7%) | 15 (26.3%) |
| Missing | 6 (5.0%) | 1 (1.6%) | 5 (8.8%) |
| Normal | 74 (62.2%) | 37 (59.7%) | 37 (64.9%) |
| SLEDAI Score* |  |  |  |
| High | 13 (10.9%) | 6 (9.7%) | 7 (12.3%) |
| Low | 106 (89.1%) | 56 (90.3%) | 50 (87.7%) |
| Anti-Sm Ab |  |  |  |
| Abnormal | 26 (21.8%) | 17 (27.4%) | 9 (15.8%) |
| Missing | 2 (1.7%) | 0 (0%) | 2 (3.5%) |
| Normal | 91 (76.5%) | 45 (72.6%) | 46 (80.7%) |
| High dsDNA Ab at blood draw |  |  |  |
| Abnormal | 50 (42.0%) | 31 (50.0%) | 19 (33.3%) |
| Normal | 69 (58.0%) | 31 (50.0%) | 38 (66.7%) |
| Proteinuria at blood draw |  |  |  |
| Abnormal | 5 (4.2%) | 3 (4.8%) | 2 (3.5%) |
| Normal | 114 (95.8%) | 59 (95.2%) | 55 (96.5%) |
| Photosensitivity |  |  |  |
| 0 | 80 (67.2%) | 40 (64.5%) | 40 (70.2%) |
| 1 | 39 (32.8%) | 22 (35.5%) | 17 (29.8%) |

### Association Between TEs and SLE Phenotypes is Cell Type Specific

We carried out a comprehensive genome wide analysis of TEs in CD4+ T cells, monocytes, B cells, and NK cells in relation to SLE subphenotypes(history of anti-SM antibody, history of anti-RNP antibody, anti-dsDNA antibody at blood draw, severe disease 2 vs mild disease, severe disease 1 vs mild disease, severe disease 2 vs severe disease 1, disease activity characterized by the SLEDAI score, IFN score, proteinuria and photosensitivity). Differential expression analysis of SLE subphenotypes identified significant (adj p-value < 0.05) TEs in each cell type and SLE subphenotype (Table 2). In CD4+ T cells there were between 3 and 38 differentially expressed elements, in monocytes cells there were between 3 and 34 differentially expressed elements, in B cells there were between 2 and 57 differentially expressed elements, and in NK cells there were between 2 and 37 differentially expressed elements (Table 2). These include two differentially expressed TEs (HML2_8p23.1b and ERVLB4_8p23.1o) in CD4+ T cells at a previously identified susceptibility locus of SLE (8p23)(21).

**Table 2.** Number of TEs Differentially Expressed with SLE phenotypes (adj pval < 0.05) adjusting for sex, age, sequencing lane, medications, and genetic PCs 1-10. (Up-regulated/Down-regulated)

|  | CD4 | CD14 | CD19 | NK |
| --- | --- | --- | --- | --- |
| Autoantibody Production |  |  |  |  |
| SM + vs - | 16 (7,9) | 3 (0,3) | 33 (6,27) | 14 (6,8) |
| RNP + vs - | 16 (6,10) | 3 (2,1) | 18 (11,7) | 16 (7,9) |
| dsDNA + vs - | 11 (4,7) | 11 (5,6) | 17 (8,9) | 19 (7,12) |
| Disease Severity |  |  |  |  |
| SD2 vs M | 32 (2,30) | 16 (0,16) | 34 (21,13) | 16 (3,13) |
| SD1 vs M | 8 (3,5) | 11 (4,7) | 20 (11,9) | 28 (9,19) |
| SD2 vs SD1 | 38 (3,35) | 34 (4,30) | 57 (38,19) | 37 (2,35) |
| SLEDAI Score High vs Low | 8 (2,6) | 23 (3,20) | 24 (1,23) | 8 (0,8) |
| Clinical Manifestations |  |  |  |  |
| IFN High vs Low | 15 (6,9) | 18 (8,10) | 7 (1,6) | 22 (0,22) |
| Proteinuria + vs - | 3 (0,3) | 13 (1,12) | 2 (1,1) | 2 (1,1) |
| Photosensitivity + vs - | 11 (8,3) | 12 (9,3) | 33 (3,30) | 22 (3,19) |

We then explored whether these differentially expressed TEs were cell or SLE subphenotype specific. No locus specific TEs were shared across all SLE subphenotypes across all cell types (CD4+ T cells, monocytes, B cells, and NK cells). Very few overlaps were observed across 2-3 cell types. For example, between CD4+ T cells, monocytes, and B cells L1FLnI_21q22.3a was differentially expressed.

We observe very few overlaps of TEs across all SLE subphenotypes and cell types (Figure 2). In CD4+ T cells we identify 1 TE (ERVLB4_8p23.1o) that was common across 5 subphenotypes including (photosensitivity, anti-dsDNA antibody at time of blood draw, disease activity characterized by the SLEDAI score, history of anti-RNP antibody, and cluster severe disease 1 vs mild disease). In the monocyte analysis, we found 2 overlapping TEs (L1FLnI_5q35.1e and L1FLnI_Yp11.2na) common across 6 SLE subphenotypes (photosensitivity, anti-dsDNA antibody at the time of blood draw, disease activity characterized by the SLEDAI score, history of anti-RNP antibody, clinical cluster severe disease 1 vs mild disease and Interferon signature). In B cells we identify 1 TE (HML2_8p23.1a) that is differentially expressed across 8 subphenotypes (photosensitivity, anti-dsdna antibody at the time of blood draw, disease activity characterized by the SLEDAI score, history of anti-RNP antibody, severe disease 2 vs severe disease 1, severe disease 1 vs mild disease, IFN signature, and anti-SM antibody). In NK cells we found 1 TE common (LTR25_9q13a) across 4 SLE subphenotypes (anti-dsdna antibody at the time of blood draw, history of anti-RNP antibody, IFN signature and anti-SM antibody).

**Figure 2:**
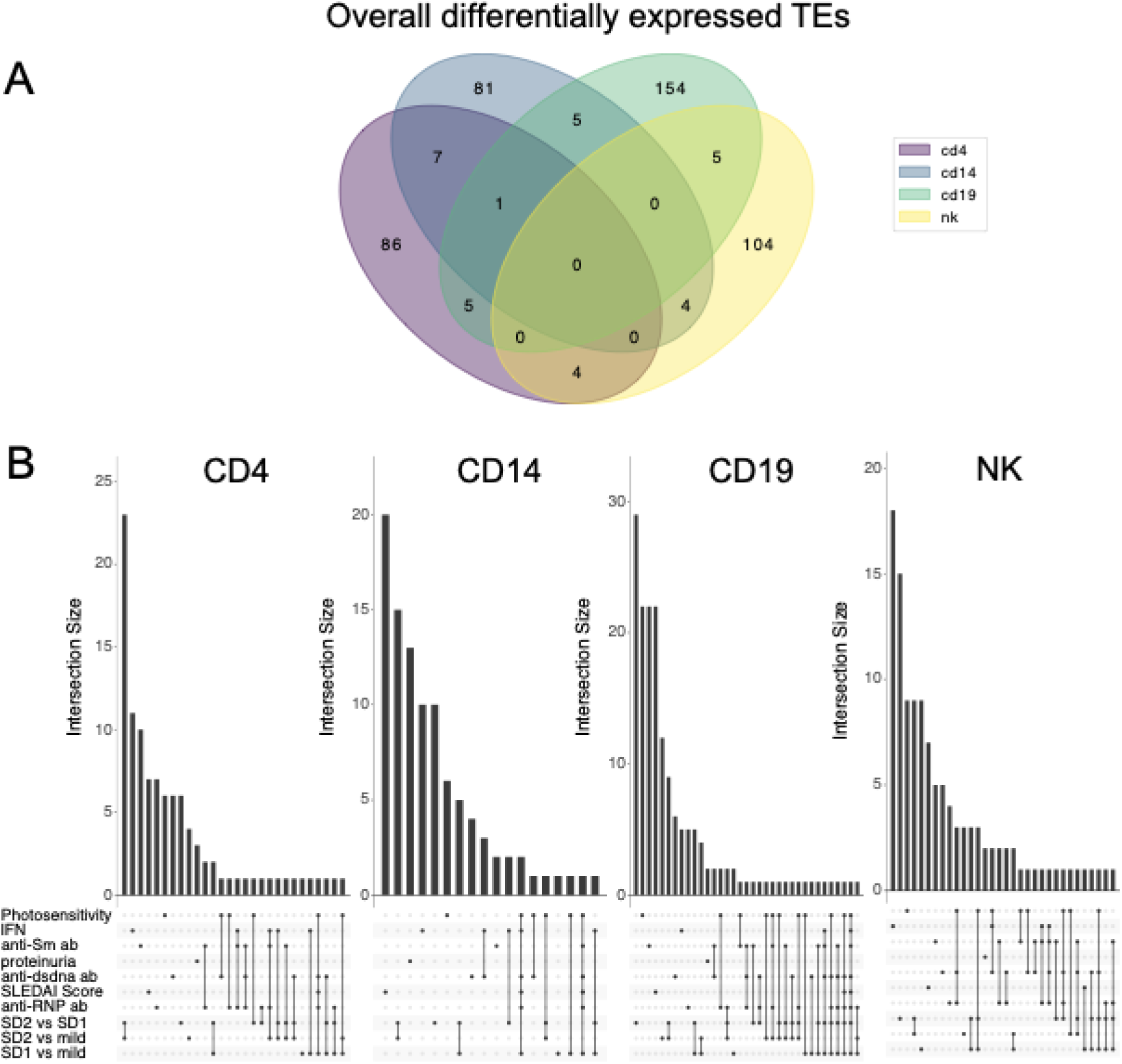
Identification of common differentially expressed TE’s in the combined cohort patients (ancestry stratified) shows distinct locus specific TEs differentially expressed across cell types and SLE subphenotypes. A. Venn diagram of overlap of DE TEs between cell types B. Overlap of DE TEs between lupus subphenotypes

### Ancestry Stratified Analysis Reveals Additional Cell-Type-Specific Associations of TEs with SLE Subphenotypes

Comparing patients based on genetic similarity revealed two strong clusters (Figure S1), which significantly correlated with self reported race (r^2^ 0.938, p-value 2.2e-16). Given the differences in SLE burden between individuals with Asian and European ancestry, we explored differential TE expression within these groups(22). Similar to our previous analysis, we identified many differentially expressed locus specific TEs by SLE subphenotype and cell type when stratifying by genetic ancestry groups and removing admixed individuals (Table 3).

**Table 3.** Stratified analysis by genetic similarity (Fig S1) of differentially expressed TEs with SLE phenotypes (adj pval < 0.05). Adjusting for sex, age, sequencing lane, medications, and genetic ancestry pcs 1-10 (Up-regulated/Down-regulated)

|  | CD4 | CD14 | CD19 | NK |
| --- | --- | --- | --- | --- |
| Autoantibody Production |  |  |  |  |
| SM + vs - | 14 (8,6) | 0 (0,0) | 0 (0,0) | 6 (1,5) |
| RNP + vs - | 158 (40,118) | 47 (12,35) | 80 (17,63) | 22 (13,19) |
| dsDNA + vs - | 107 (26,81) | 24 (9,15) | 167 (44,123) | 30 (6,24) |
| Disease Severity |  |  |  |  |
| SD2 vs M | 40 (15,25) | 0 (0,0) | 22 (16,6) | 4 (2,2) |
| SD1 vs M | 123 (45,78) | 50 (21,29) | 113 (29,84) | 28 (9,19) |
| SD2 vs SD1 | 55 (20,35) | 0 (0,0) | 54 (38,16) | 8 (5,3) |
| SLEDAI Score High vs Low | 28 (17,11) | 14 (13,1) | 23 (11,12) | 3 (2,1) |
| Clinical Manifestations |  |  |  |  |
| IFN High vs Low | 6 (4,2) | 28 (11,17) | 0 (0,0) | 0 (0,0) |
| Proteinuria + vs - | 0 (0,0) | 0 (0,0) | 0 (0,0) | 0 (0,0) |
| Photosensitivity + vs - | 142 (28,114) | 53 (9,44) | 142 (54,88) | 68 (12,56) |

Participants self-reported as Asian and clustered by Genetic PC1
|  | CD4 | CD14 | CD19 | NK |
| --- | --- | --- | --- | --- |
| Autoantibody Production |  |  |  |  |
| SM + vs - | 102 (26,76) | 62 (40,22) | 105 (69,36) | 92 (42,50) |
| RNP + vs - | 193 (108,85) | 85 (43,42) | 123 (81,42) | 203 (128,75) |
| dsDNA + vs - | 229 (133,96) | 113 (75,38) | 228 (157,71) | 52 (27,25) |
| Disease Severity |  |  |  |  |
| SD2 vs M | 40 (12,28) | 26 (6,20) | 84 (69,15) | 137 (114,23) |
| SD1 vs M | 122 (82,40) | 74 (46,28) | 147 (61,86) | 270 (45,225) |
| SD2 vs SD1 | 201 (36,165) | 115 (27,88) | 235 (136,99) | 162 (108,54) |
| SLEDAI Score High vs Low | 42 (28,14) | 19 (13,6) | 35 (27,8) | 54 (47,7) |
| Clinical Manifestations |  |  |  |  |
| IFN High vs Low | 139 (86,53) | 113 (77,36) | 54 (37,17) | 319 (203,116) |
| Proteinuria + vs - | 1 (1,0) | 1 (0,1) | 0 (0,0) | 0 (0,0) |
| Photosensitivity + vs - | 240 (48,192) | 143 (38,105) | 291 (56,235) | 303 (29,274) |

We observe a larger number of differentially expressed TEs in lupus subphenotypes, especially anti-dsDNA in this stratified analysis when compared to the analysis with all patients (Table 3). As in the overall analyses, few locus-specific TEs were differentially expressed across cell types or subphenotypes in the stratified analyses (Figures 3 and 4).

**Figure 3.**
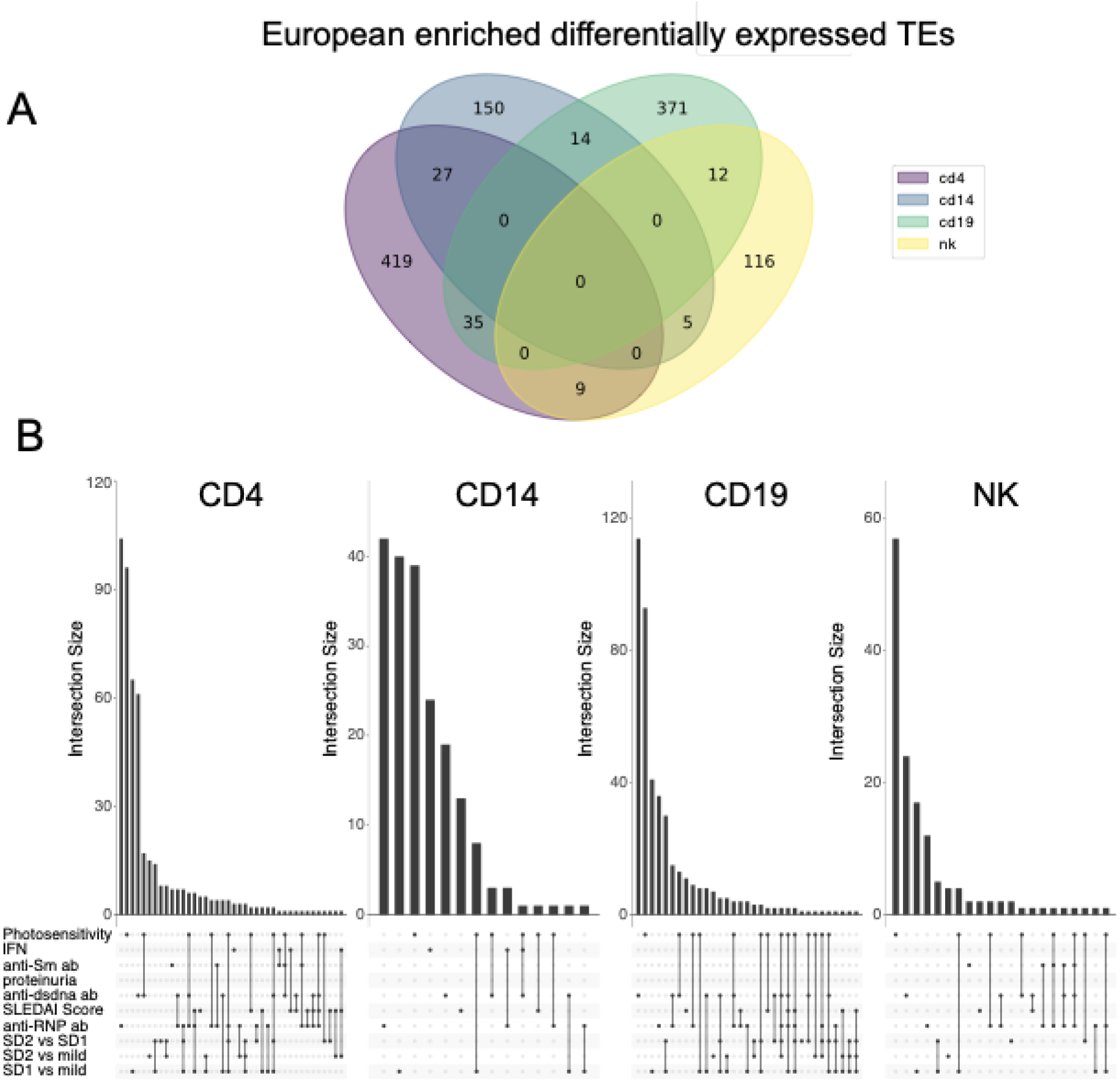
Identification of common differentially expressed TE’s in the European enriched cohort patients (ancestry stratified) shows distinct locus specific TEs differentially expressed A. across cell types and B. SLE subphenotypes

**Figure 4.**
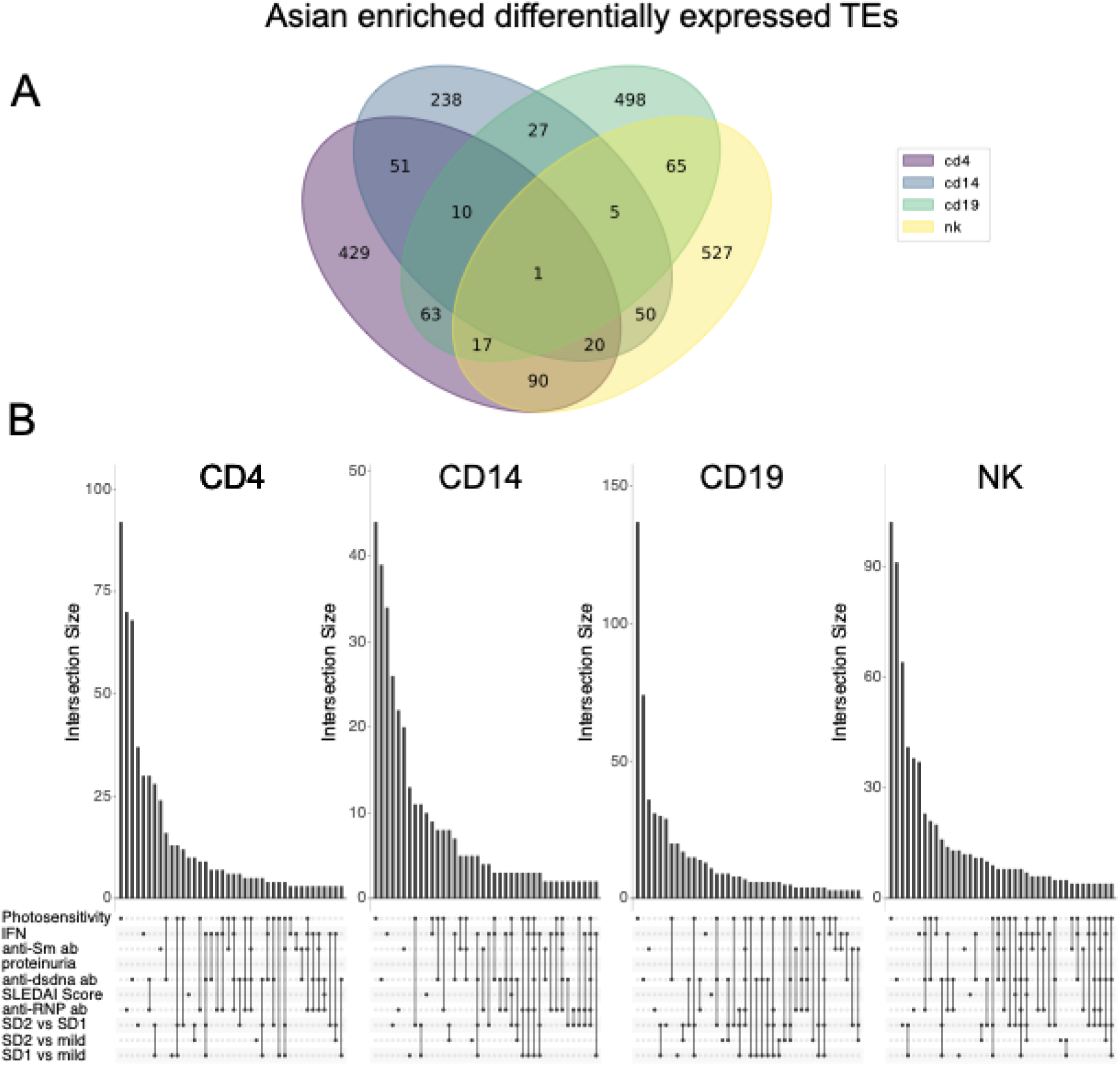
Identification of common differentially expressed TE’s in the Asian enriched cohort patients (ancestry stratified) shows distinct locus specific TEs differentially expressed A. across cell types and B. SLE subphenotypes

In the Asian enriched subgroup we saw the largest number of overlaps across cell types. There were 10 TEs in common across CD4+ T cells and monocytes, 5 TEs common between monocytes, B cells, and NK cells, 20 TEs common between CD4+ T cells, monocytes, and NK cells and 5 TEs common between monocytes, B cells, and NK cells. The European enriched subgroup had fewer TEs shared across phenotypes and cell types in comparison to the Asian enriched subgroup.

In the European enriched subgroup of patients in CD4+ T cells we found 1 TE (MER101_21q21.3a) common across 4 different subphenotypes (anti-dsDNA antibody, photosensitivity, disease activity characterized by the SLEDAI score, and Anti-RNP antibody). In monocytes we identified 1 TE (HERVL_Xp11.4) common across three SLE subphenotypes (photosensitivity, anti-dsDNA antibody and IFN). In B cells we found two TE’s (ERVLB4_2q11.1b, HARLEQUIN_Yq11.223) that are shared across 5 SLE subphenotypes (photosensitivity, anti-dsDNA antibody, disease activity characterized by the SLEDAI score, anti-RNP antibody, and severe disease 1 vs mild disease). In NK cells we found one TE (L1FLnI_8q13.1d) common across four SLE subphenotypes (photosensitivity, anti-SM antibody, anti-RNP antibody, and anti-dsDNA).

In the Asian ancestry subgroup of patients a in CD4+ T cells there were 3 common differentially expressed TEs (ERVL_17q11.1, LTR23_3q26.31, and MER41_17q23.3a) across 4 SLE subphenotypes (anti-dsDNA antibody, anti-RNP antibody, IFN, anti-SM antibody). In monocytes cells there were 2 TEs (L1FLnI_6q14.3u, MER101_6p21.1) across 6 subphenotypes (anti-dsDNA antibody, anti-RNP antibody, severe disease 2 vs severe disease 1, Anti-SM antibody, photosensitivity, and IFN). In B cells there were 4 differentially expressed TEs (ERV316A3_4q28.3cl, ERVLE_4q32.3a, MER4B_15q21.1c, MER4B_Xq22.3a) across 4 SLE subphenotypes (photosensitivity, anti-dsDNA antibody, severe disease 2 vs severe disease 1, and severe disease 1 vs Mild Disease). Lastly, in NK cells there were 4 common differentially expressed TEs (ERVLB4_Xq21.31j, HUERSP2_Xq27.3a, L1FLnI_11p14.3k, LTR25_16p12.3b) across four SLE subphenotypes (photosensitivity, anti-dsDNA antibody, anti-RNP antibody, and severe disease 2 vs severe disease 1)

### Family-level enrichment of TEs and TE derived viral gene analysis shed light on differentially expressed TEs across diverse SLE subphenotypes within each unique cell type

Most families of TEs have deposited sequences at thousands of loci throughout the genome. Given their common origin, these locus-specific instances of each TE share very similar sequences and potential functional elements. Thus, expression of similar sequences from different genomic loci derived from the same family of TEs might collectively contribute to SLE. Previous studies have associated HERV families, like HERV-K, with SLE(20).

To test for family-level effects, we analyzed differentially expressed TEs at the family level across subphenotypes of SLE, focusing on each cell type. We discovered significant enrichment of different families among the differentially expressed TEs associated with SLE sub-phenotypes in each cell type (Figure 5A). In CD4+ T cells, we observed enrichment of HERVH (adj p-value 1.35E-29), while in monocytes, we detected enrichment for MER61(adj p-value 0.01). In B cells, we found HML2 (adj p-value 1.47E-04) and HERVH (adj p-value 1.07E-30) to be enriched, and in NK cells ERVLB4 (adj p-value 1.53E-06) was enriched. We also discovered consistent depletion for L1FLnI sequences across all four cell types.

**Figure 5.**
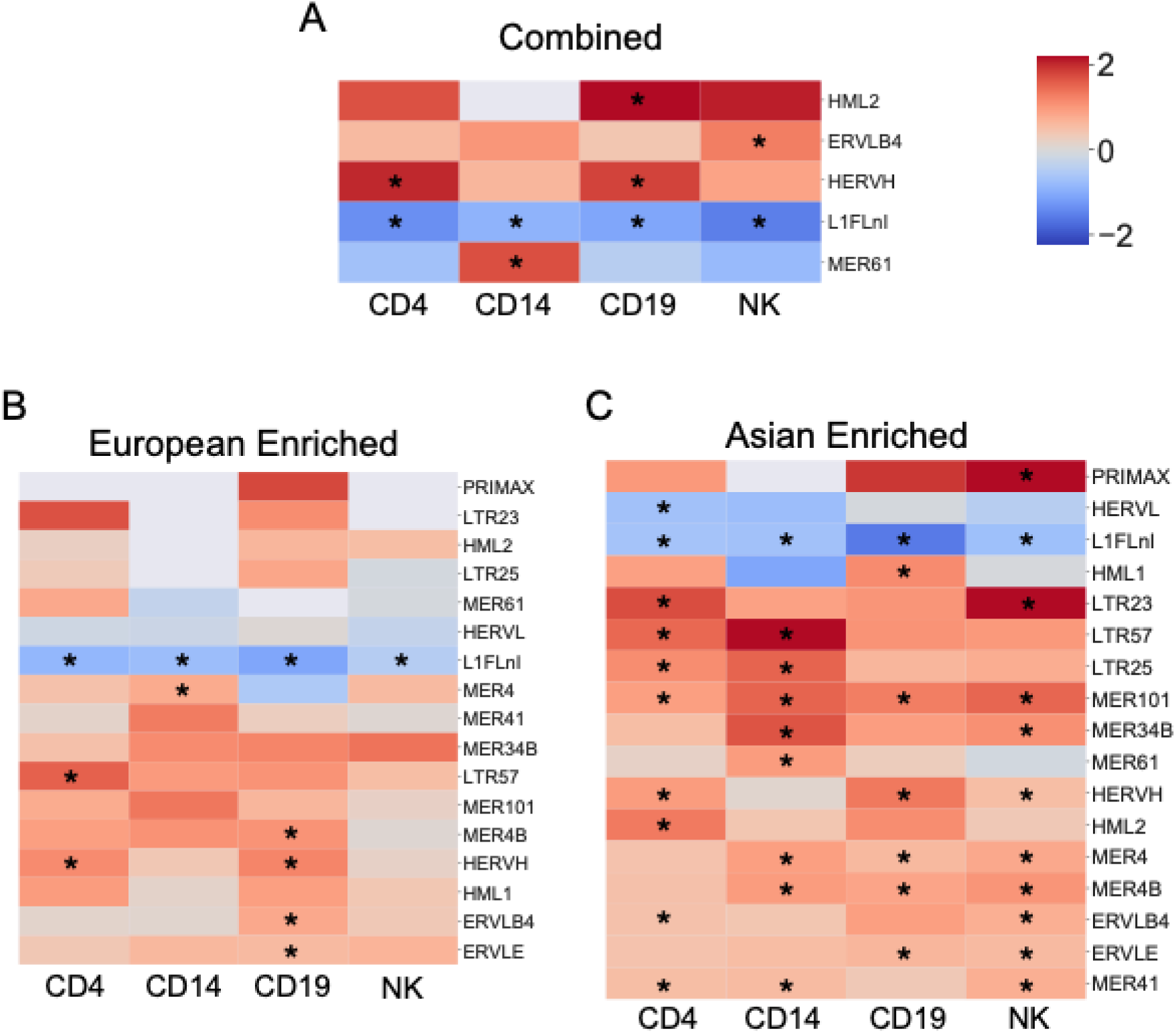
Log odds ratio of significant enrichment/depletion of HERV families in combined and stratified analysis A. Family enrichment for the combined analysis B. Family enrichment for the European enriched cohort C. Family Enrichment for the Asian enriched cohort. Family annotation for HERV families used from https://github.com/mlbendall/telescope_annotation_db (significant families denoted by *)

In the analysis stratified by genetic similarity, notable distinctions emerged. In the Asian enriched group, TEs manifested unique patterns of enrichment across different cell types. Specifically, MER101 was consistently enriched across all cell types. NK cells in this subgroup were characterized by the enrichment of PRIMAX, LTR23, MER34B, HERVH, MER4, MER4B, ERVLB4, ERVLE, and MER41. Meanwhile, B cells displayed enrichment of HML1 and HERVH, MER4, MER4B, ERVLE; and monocytes cells showed enrichment of LTR57, LTR25, MER34B, MER61, MER4, MER4B, and MER41. Enrichment in CD4+ T cells was observed for LTR23, LTR57, LTR25, HERVH, HML2, ERVLB4, and MER41; while L1FLnI demonstrated consistent depletion across all cell types.

In comparison, in the enriched European group, HERVH and LTR57 were enriched in CD4+ T cells, whereas monocytes cells displayed enrichment of MER4. MER4B, HERVH, ERVLB4, ERVLE enrichment was characteristic of B cells. As seen in the Asian enriched group, L1FLnI showed depletion across all cell types as well.

Previous work has hypothesized that HERVs with preserved open reading frames could produce proteins which could activate or depress the inflammatory cascade(23). Therefore we used gEVE: a genome-based endogenous viral element database to find open reading frames of viral proteins derived from HERVs in the differentially expressed TEs we observed across SLE subphenotypes. We found enrichment of open reading frames of viral proteins derived from TEs in the cell specific differentially expressed TEs across all SLE subphenotypes (Table 4).

**Table 4.** Viral protein enrichment using gEVE (a genome-based endogenous viral element database) gag-structural components of matrix, capsid, and nucleocapsid, pro - protease, RT-reverse transcriptase, int - integrase, env - envelope-associated glycoprotein, dut- dUTPase ^32^

|  | CD4 | CD14 | CD19 | NK |
| --- | --- | --- | --- | --- |
| Combined | pro, int, RT |  | pro, RT | pro, env, RT |
| European Enriched | pro, int, gag, dut, RT |  | pro, int, RT |  |
| Asian Enriched | pro, RT | pro, dut, int, RT | pro, int, RT | pro, RT |

### Interferon Pathway Associated with Expression of Differentially Expressed TEs

To better understand possible effects of differential TE expression in SLE, we asked whether the expression of host genes correlated with the expression of TEs associated with SLE subphenotypes in our cohort. The expression levels of many genes associated with expression of DE TEs in the combined analysis (Table 5, Figure 6). The associations of differentially up and down regulated genes were largely cell-type-specific (Figure 6E and 6F). In the CD4+ T cells there is a strong upregulation of genes associated with the differentially expressed TEs. Some of the strongest up regulated genes in CD4+ T cells were involved in type I interferon signaling and other antiviral innate immune pathways such as *LY6E, IFI6, ISG15, and ISG20*. In the monocyte analysis, most of the genes also appear upregulated as seen in the volcano plots, and the top upregulated genes are also involved in antiviral activity such as *ISG15, IFI6, IFI35, BST2, and TRIM22*. In CD19 cells there are also many up-regulated genes as seen in the volcano plot. Some of the top upregulated genes are *ISG15, IFIT3, IRF7, and BST2* which play roles in response to viruses. In NK cells there are many up and down regulated genes as seen in 6D, some of the top upregulated genes are *NXF1*, which is involved in transport of unspliced retroviral genomic RNA and *CAPZA1* which was downregulated, and is also observed to be downregulated in virus infected cells(24, 25).

**Figure 6.**
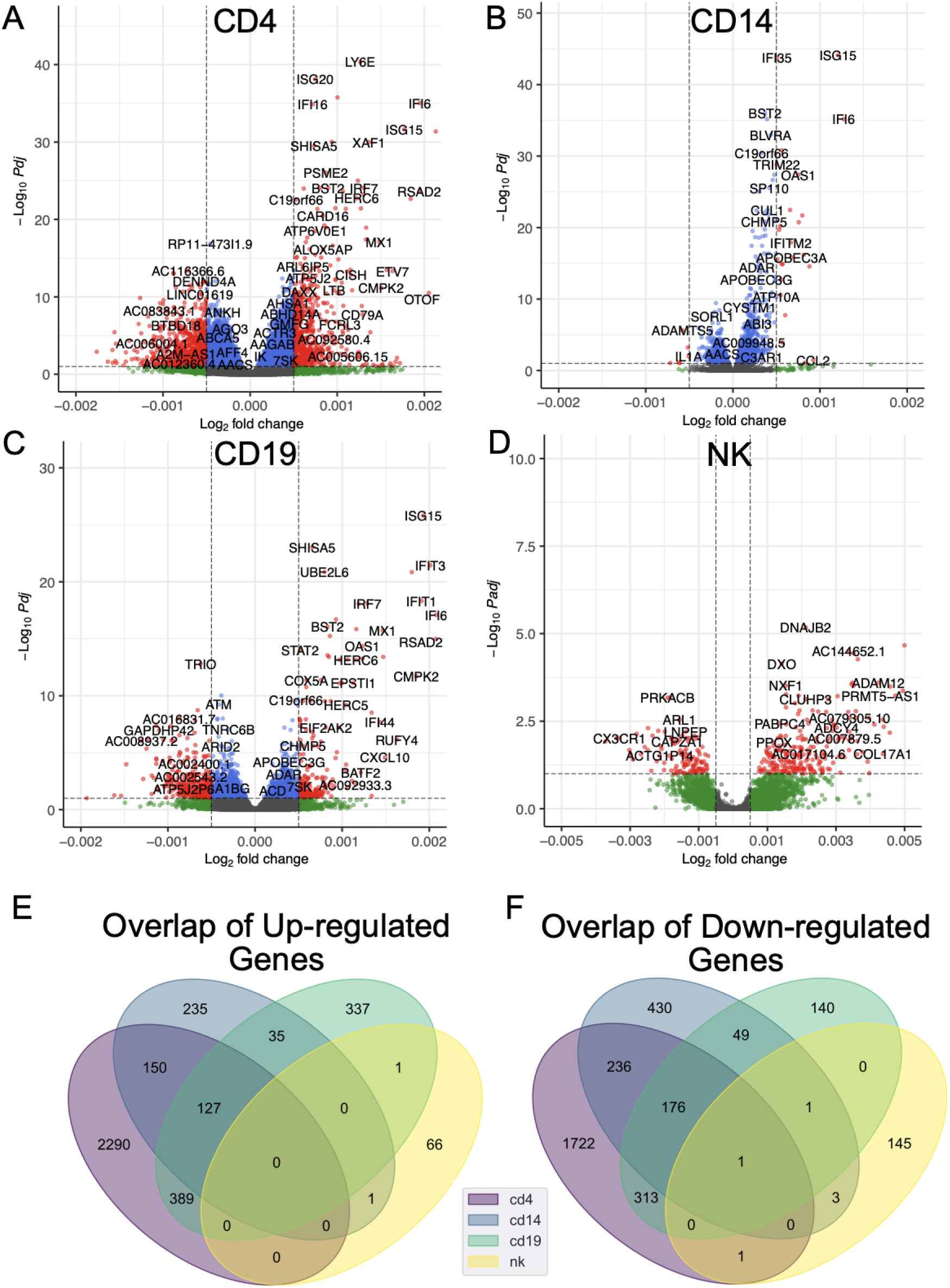
Cell specific volcano plots of combined analysis differentially expressed gene associations with differential TE expression using DESeq2 A. CD4, B. CD14, C. CD19 and D. NK. Identification of the common DE genes (padj < 0.05) across the different cell types: E. up-regulated F. down-regulated

**Table 5.** Differential Gene Expression with respect to lupus associated TEs (Up-regulated/Down-regulated)

|  | CD4 | CD14 | CD19 | NK |
| --- | --- | --- | --- | --- |
| Combined | 5405 (2449,2956) | 1444 (896, 548) | 1569 (680,889) | 219 (151,68) |
| European Enriched | 34 (5,29) | 223 (115, 108) | 70 (9,61) | 288 (221,67) |
| Asian Enriched | 18 (9,9) | 17 (3,14) | 16 (0,16) | 24 (5,19) |

Gene set enrichment analysis revealed that SLE associated TEs correlated with differential expression of similar pathways, such as interferon and cytokine signaling, across cell types. Pathways expressed in all cell types except NK cells include Interferon alpha/beta signaling, Interferon Signaling, Cytokine Signaling In Immune system, and antiviral mechanism by IFN-stimulated genes(Figure 7). There are also some cell type specific signals that were identified. For example, differentially enriched pathways with TE expression include SLC-mediated transmembrane transport in monocytes cells; influenza infection, viral mRNA translation, antigen presentation: folding assembly and peptide loading of class I MHC in CD4 cells; GPCR ligand binding, host interactions with influenza factors, and NS1 mediated effects on host pathways, in B cells(Supplementary Tables 6).

**Figure 7.**
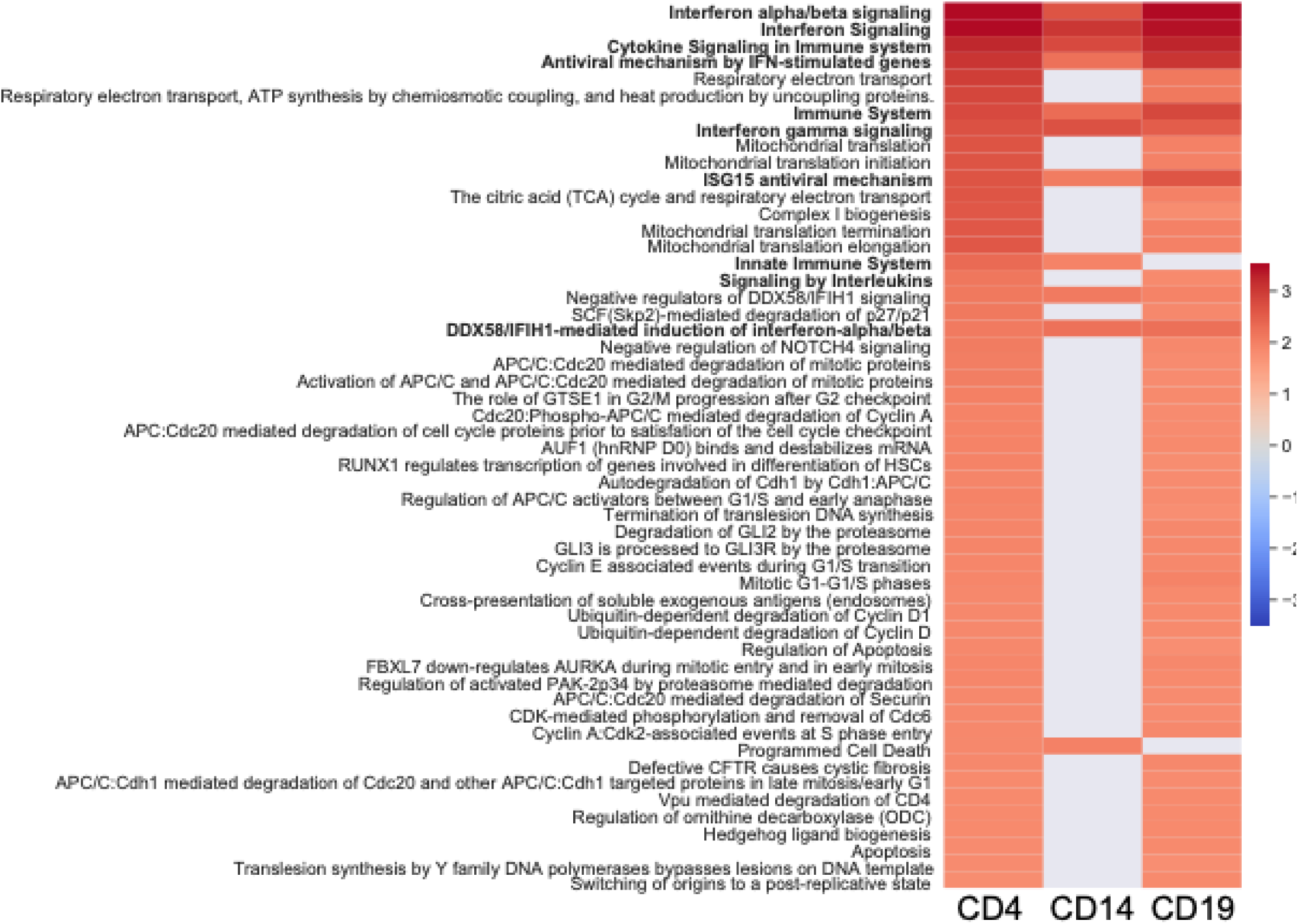
Significant pathways in more than one cell type from gene set enrichment analyses with webgestalt using significant genes in the combined cohort. Heatmap shoes normalized enrichment score

## Discussion

This is the first study to characterize locus and cell-specific TE expression among a deeply phenotyped SLE cohort. We found that TE differential expression is highly cell specific, with very few overlaps between immune cells. We discovered associations with several disease subphenotypes and host gene expression. This suggests that HERV expression could contribute to the heterogeneity of SLE across individuals and populations.

We found independent associations of TE expression with production of autoantibodies against ribonucleoproteins (RNP, SM), SLE disease activity and SLE disease severity, after adjusting for potential confounders. Some of these results have been previously reported; for example, some TEs, such as HERV-E clone 4-1 have correlated with autoantibody levels (anti-U1 RNP, anti-SM nuclear antibodies) and higher interferon status in SLE(26, 16). These associations are highly cell specific, subphenotype specific, which suggests that their differential expression could contribute or be a result of different lupus subphenotypes.

Given the differences in SLE severity and outcomes between patients who self-report as Asian or White, and that TE are encoded through the germline, we explored TE differential expression patterns in individuals stratified by genetic similarity. The genetic similarity analysis showed a near complete agreement between genetic similarity based clustering and self-reported race in this cohort, as well as a small number of individuals with significant genetic admixture. Admix individuals were removed from downstream stratified analyses (Figure S1). When the two groups, Asian enriched and European enriched were stratified, we found more differentially expressed TEs in almost every SLE subphenotype, despite adjusting for genetic principal components 1-10. This was surprising given that the sample size for these groups was reduced from the combined analysis. Further work is needed to determine if these differences in the stratified analysis between TE expression and phenotypes are due to genetic factors or due to differences in environmental exposures.

Retroviral p30 gag proteins and serum reactivities to p30 gag antigen have been found in SLE patients with proliferative glomerulonephritis(27). We investigated if it was possible that the differentially expressed TEs identified exert their effect as proteins, as most expressed TEs do not contain full open reading frames for functional viral proteins. We tested if there was significant enrichment of open reading frames among the identified differentially expressed TEs and found enrichment of several TE derived viral proteins in a cell specific manner, such as pro, env, and RT. The role of TE derived viral proteins in SLE remains to be elucidated, however previous work has found env proteins both to trigger innate and adaptive immunity as well as possessing immunosuppressive properties(28). Therefore it is difficult to say whether the enrichment of viral proteins we observe in the differentially expressed TEs is the cause of an immune response, or the response acting to downregulate an immune response.

One of the most intriguing findings was the connection between host response gene expression and the expression of transposable elements (TEs) in relation to the SLE phenotypes. It has been hypothesized that TEs could activate the innate immune system, and unlike a viral infection where the immune response is curtailed, could lead to chronic inflammation and contribute to autoimmunity(14). For example, the chronic induction of type I IFNs which could put cells into an antiviral state, shutting down metabolic processes and leading to apoptosis(14). Supporting this, a study of kidney biopsies from SLE patients with lupus nephritis found LINE elements can trigger the IFN-I pathway(29). We found that differentially expressed TEs are associated with genes and pathways that are involved in an antiviral response, which lends support to this hypothesis. There are also cell specific pathways of note, for instance, in CD4+ T cells, we observe a correlation between differentially expressed TEs and many pathways indicating response to viral infections, which have long been associated with SLE, such as influenza infection and life cycle, viral mRNA translation, influenza viral RNA transcription and replication. In B cells, we identify associations with inflammatory pathways, including NF-kappaB, which have been associated with both lupus and viral infection(30, 31). Additional studies are needed to examine causality between TE expression and immune dysregulation in order to understand whether TE expression contributes to the development of SLE.

Nonetheless, there exist several limitations within this study. The absence of healthy control data hampers a cell-specific comparison to a baseline TE expression profile and the total sample size is limited. While the cohort described here has individuals with Asian and European ancestry, it would be interesting if these findings hold up in a more diverse population from the US and beyond. Furthermore, most of the participants in this cohort exhibit low disease activity, thereby limiting our power to detect DE TEs across different levels of disease activity. Finally, though we were able to account for important potential confounding factors such as medication usage, age, and gender, we cannot exclude the possibility of additional unidentified confounders on TE expression. As this study design is cross-sectional, it is difficult to ascertain whether TE expression is a consequence, rather than causal in the mechanism of SLE manifestations, and further studies are needed to understand these results.

This study aimed to investigate the impact of TE expression on SLE heterogeneity, in a cell-specific manner. We discover a significant number of locus-specific TEs and TE families whose expression is associated with specific SLE manifestations and host gene expression. In summary, our study reveals cell-specific TE expression patterns linked to disease activity, autoantibody production, and distinct disease manifestations such as lupus nephritis. TE expression is associated with expression of host genes that are relevant to SLE pathogenesis. To gain deeper insights, further exploration into TE expression within normal immune cells might be needed. In addition, further functional studies could potentially shed light on whether TE expression contributes to the pathogenesis of SLE.

## Methods

### Cohort Description and Data Generation

All patients in this study were from the California Lupus Epidemiology Study (CLUES), a cohort of individuals with physician-confirmed SLE. CLUES was approved by the Institutional Review Board of the University of California, San Francisco. All participants signed a written informed consent to participate in the study. Study procedures involved an in-person research clinic visit, which included collection and review of medical records prior to the visit; history and physical examination conducted by a physician specializing in lupus; a collection of biospecimens, including peripheral blood for clinical and research purposes; and completion of a structured interview administered by an experienced research assistant. All SLE diagnoses were confirmed by study physicians based upon one of the following definitions: (a) meeting ≥4 of the 11 American College of Rheumatology (ACR) revised criteria for the classification of SLE as defined in 1982 and updated in 1997, (b) meeting 3 of the 11 ACR criteria plus a documented rheumatologist’s diagnosis of SLE, or (c) a confirmed diagnosis of lupus nephritis, defined as fulfilling the ACR renal classification criterion (>0.5 grams of proteinuria per day or 3+ protein on urine dipstick analysis) or having evidence of lupus nephritis on kidney biopsy. Based on sample availability at the time of sequencing, a total of 120 patients (Supplementary Data 1) were profiled with bulk RNA-seq from the CLUES cohort. Clinical data collected at sampling and the self-reported race was used for downstream analyses.

#### Subphenotype Definitions

Disease activity was measured with the standardized disease activity score SLEDAI(32). A high SLEDAI score was defined as a score greater than or equal to 8, while low was defined as less than 8. Besides the total SLEDAI score, we also performed analyses with specific items of the SLEDAI such as proteinuria and presence of dsDNA antibody. From our own previous work, three stable clusters named mild disease, severe disease 1, and severe disease 2, were revealed from unsupervised clustering of the 11 American College of Rheumatology (ACR) Classification Criteria characterized by significant differences in SLE manifestations(33). We also performed subphenotype associations with history of presence of anti Smith antibody (anti-Sm), anti ribonucleoprotein antibody (anti-RNP) and anti double stranded DNA antibody (anti dsDNA). Photosensitivity was defined as a rash or feeling sick after going out in the sun. Serologies were performed in CLIA certified labs and reported as abnormal or normal.

#### RNA-Seq Data Generation

Peripheral blood mononuclear cells were isolated from 120 SLE participants. Using the EasySep protocol from STEM cell technologies, these cells were sorted into CD14+ monocytes, B cells, CD4+ T cells, and NK cells, for a total of 480 samples. These samples were sequenced on a HiSeq4000 PE150 and gene expression data was generated using Salmon v0.8.2 with adapter-trimmed reads, quality control was performed as previously described(2).

#### Genotyping

Genotyping for genomic DNA from peripheral blood was performed using the Affymetrix Axiom Genome-Wide LAT 1 Array. This genotyping array is composed of 817,810 SNP markers across the genome and was specifically designed to provide maximal coverage for diverse racial/ethnic populations, including West Africans, Europeans and Native Americans(34). Samples were retained with Dish QC (DQC) ≥ 0.82. SNP genotypes were first filtered for high-quality cluster differentiation and 95% call rate within batches using SNPolisher. Additional quality control was performed using PLINK. SNPs having an overall call rate less than 95% or discordant calls in duplicate samples were dropped. Samples were dropped for unexpected duplicates in identity by descent (IBD) analysis or mismatched sex between genetics and self-report; for first-degree relatives, one sample was retained. All samples had at least 95% genotyping and no evidence of excess heterozygosity (maximum < 2.5*SD). We tested for Hardy-Weinberg Equilibrium (HWE) and cross-batch association for batch effects using a subset of subjects that were of European ancestry and negative for ds-DNA antibodies and renal disease to minimize genetic heterogeneity. SNPs were dropped if HWE *p* < 1e-5 or any cross-batch association *p* < 5e-5. Genetic PCs were generated using EIGENSTRAT and used for patient stratification.

### Transposable Element QC and Expression Quantification

Transposable element expression quantification was performed using adapter-trimmed reads from the RNA-Seq data that were aligned with bowtie2 to hg38 allowing for 100 alignments per read, using the very sensitive local setting (-k 100 –very-sensitivie-local –score-min L,0,1.6). Sorted bams were used as input into Telescope with default settings and the reference retro.hg38.v1 annotation from https://github.com/mlbendall/telescope_annotation_db/tree/master/builds. Locus specific TE read counts generated by Telescope were used for downstream analysis.

### Differential TE Expression Analysis

TE differential expression analysis was run with DESeq2 (v.1.38.3) using counts from Telescope with outliers dropped per cell type adjusting for age, sequencing lane, sex, genetic ancestry PCs 1-10, and medication use at the time of blood draw. For data analyses, we grouped immunosuppressive medications into the following categories: biologic treatments (belimumab, abatacept, rituximab), low-dose prednisone (<10 mg), moderate or high-dose prednisone (>10 mg), antimalarials, calcineurin inhibitors, methotrexate and leflunomide, azathioprine, mycophenolate mofetil, and cyclophosphamide. Medications were used as factors in DESeq2 as covariates. Outcomes studied included disease activity (SLEDAI score), interferon signature high/low based on gene signature developed by Kennedy et al(35), photosensitivity (previous work has found UV light induces HERV expression(36)), proteinuria, autoantibody production (dsDNA, RNP, SM), and disease severity (as defined by clinical clusters previously described in the same SLE participants)^1^. For stratified analyses, patients were stratified according to genetic similarity (PC1 >0.025 - Asian ancestry, <-0.025 - European ancestry) and admixed individuals were not considered for downstream stratified analysis. P-values from DESeq2 with a NA value were reassigned a value of 1, and all p values from the different cell type and subphenotype analysis were FDR-corrected using Benjamini-Hochberg. Differentially expressed TEs (padj < 0.05) for all SLE subphenotypes per cell type were used for downstream analysis. PCAs of TE expression were computed with the factoextra package (v 1.0.7). Cell type PCAs were made using the variance stabilizing transformation with the DESeq2 function vst(), and VST-transformed data was visualized with plotPCA in the DESeq2 package.

### Characterization of Differentially Expressed TEs

TE group and family definitions were taken from Telescope using families.tsv from https://github.com/mlbendall/telescope_annotation_db/tree/master/builds. Locus-specific differentially expressed TE’s across all SLE subphenotypes per cell type were used to calculate family enrichment. Counts per family were generated by dropping the locus from the telescope transcript name and summing the number of differentially expressed TEs according to families.tsv. Line elements were grouped into L1FLnI, L1ORF2, and L1FLI. Log odds ratio and hypergeometric test for enrichment/depletion were calculated and an expression threshold of 4 was used for filtering.

Enrichment of viral proteins was calculated using the Genome-based Endogenous Viral Element Database (gEVE). Bedtools was used to find the intersection between Hsap38.geve.v1.bed regions and Telescope annotation regions. The differentially expressed elements from the SLE subphenotype analysis that overlapped full regions of the gEVE annotation were used to calculate enrichment of viral proteins.

### TE and Gene Expression Integration

Association of differential TE expression with gene expression was calculated using the sum of counts of differentially expressed TEs for all SLE subphenotypes per cell type. Raw counts from Telescope were normalized with DESeq2 and the counts of differentially expressed TEs for each cell type were summed to get counts of differentially expressed TEs per patient for each cell type. DESeq2 (v.1.38.3) was used with the continuous variable of summed differential counts of TEs, adjusted for age, sequencing lane, sex, genetic PCs, and medication use at the time of blood draw. Gene set enrichment analysis was run with significant genes (padj < 0.05) using WebGestaltR (v 0.4.5) and the reactome database. Volcano plots were generated using Enhanced Volcano package (v 1.16.0)

### Statistics

R was used to determine statistical significance of differences and a padj value of less than 0.05 was considered significant.

### Study Approval

Participants were recruited from the California Lupus Epidemiology Study (CLUES). CLUES was approved by the Institutional Review Board of the University of California, San Francisco. All participants signed a written informed consent to participate in the study.

## Data Availability

Raw data of this study are openly available in GEO: GSE164457. All other data are available from the corresponding author upon reasonable request.

## Author Contributions

Z.C.: study design, data analysis, quality control analyses, data interpretation, and paper preparation. C.J.Y., K.E.T., L.M.: data generation, quality control analyses, genetic ancestry estimates, and paper revision. M.D., J.Y.: CLUES co-investigators. Patient enrollment and clinical, demographic characterization of patients, and paper revisions. L.A.C.: data generation, and paper revisions. JAC, CL, GF, SP: study design, data interpretation, and paper revisions. CML: Study design, paper preparation, and paper revision. M.S.: Computational expertise, study design, data interpretation, and paper revisions. All authors edited and critically revised the manuscript for important intellectual content and gave final approval for the version to be published.

## Acknowledgements

We are grateful to all patients who participated in this study. We are also grateful to Jane Hughan who provided feedback on the manuscript. This study was funded through the following grants: P30 AR070155 (M.S., Z.C.)

## Supplementary Figures

**Supplementary Figure 1.**
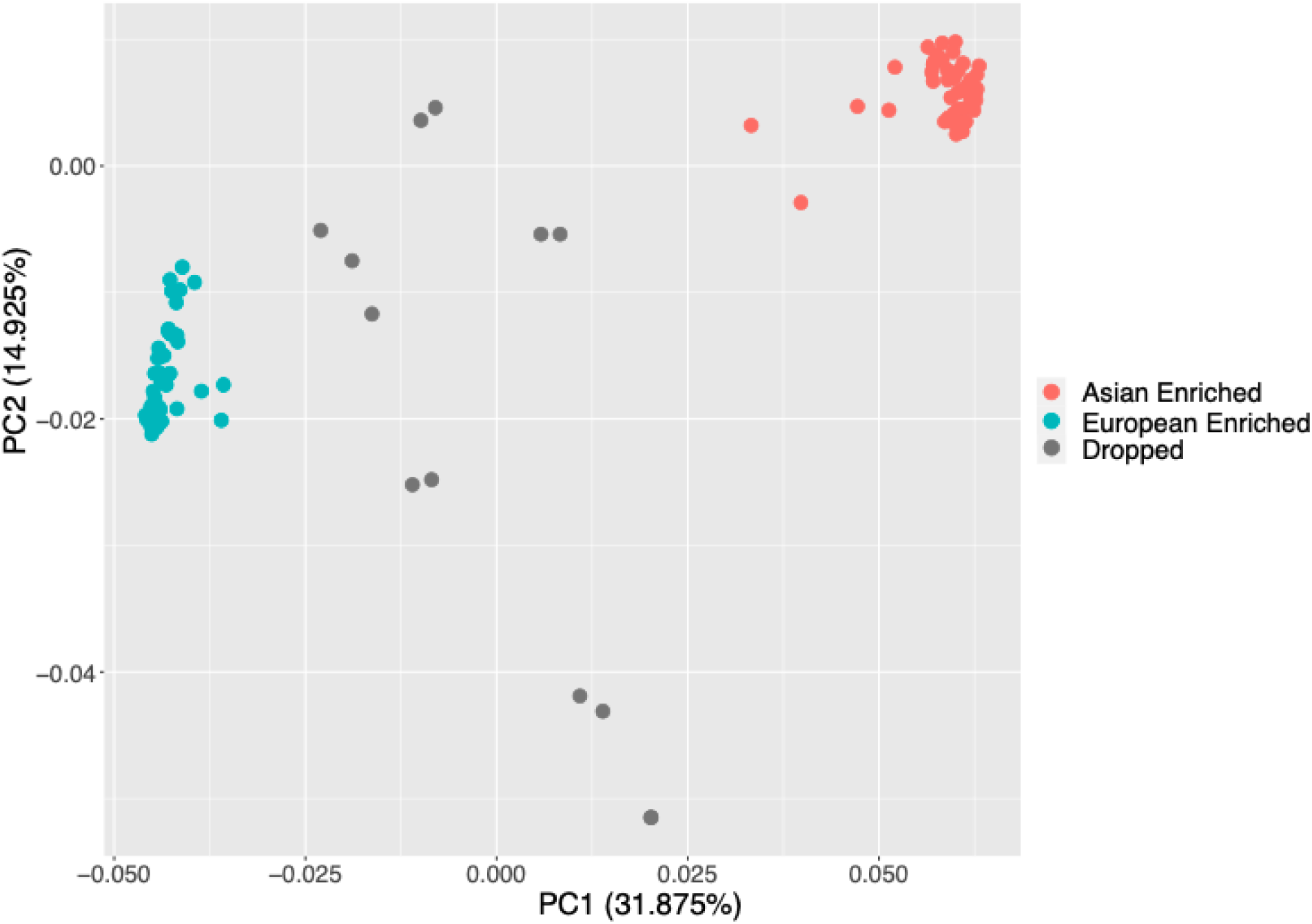
PCA plot of genotyping data across 120 patients. Individuals are colored by ancestry. Individuals in gray are considered admixed and were dropped from the stratified analysis.

**Supplementary Figure 2.**
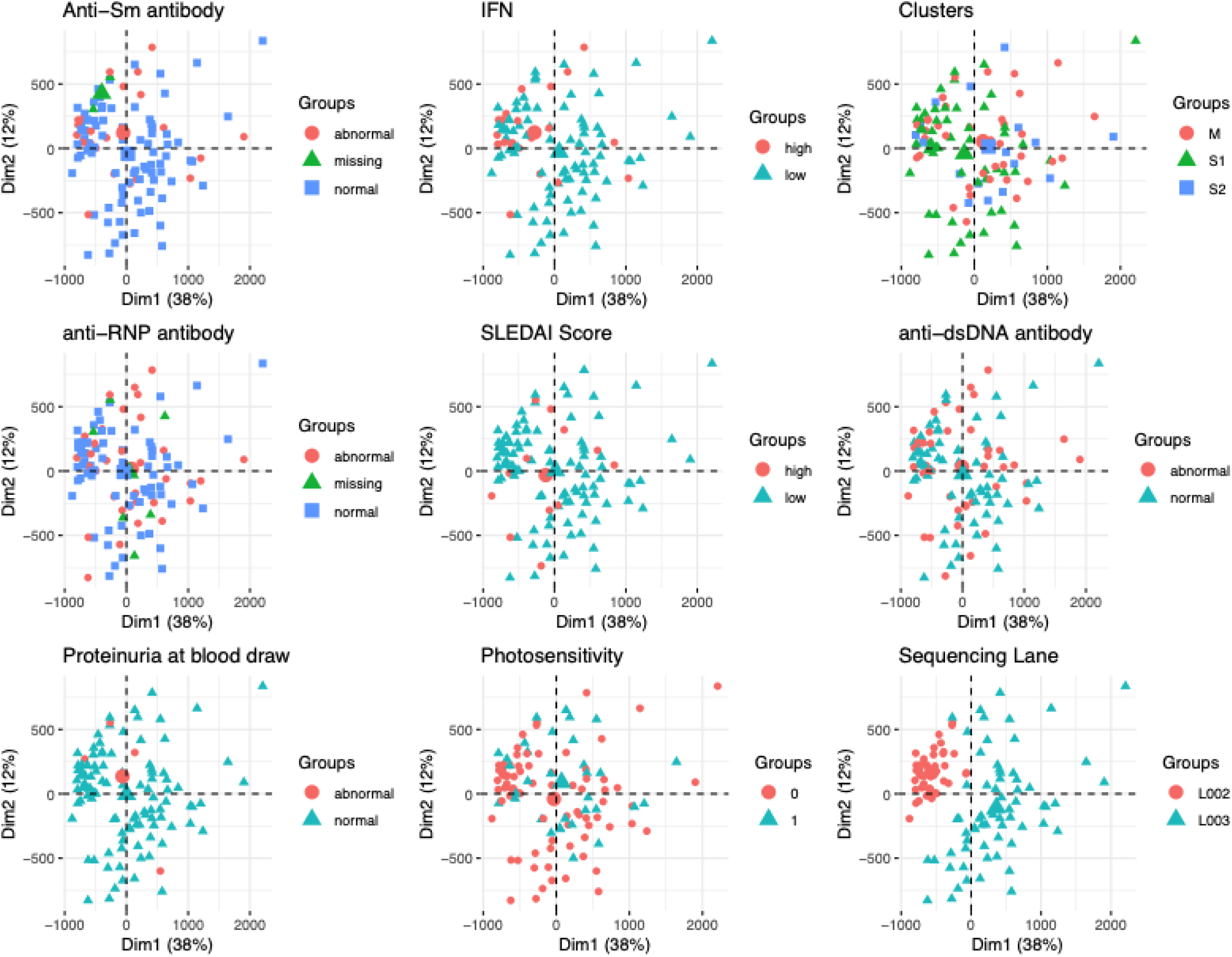
CD4 TE PCAs colored by demographics and phenotypes

**Supplementary Figure 3.**
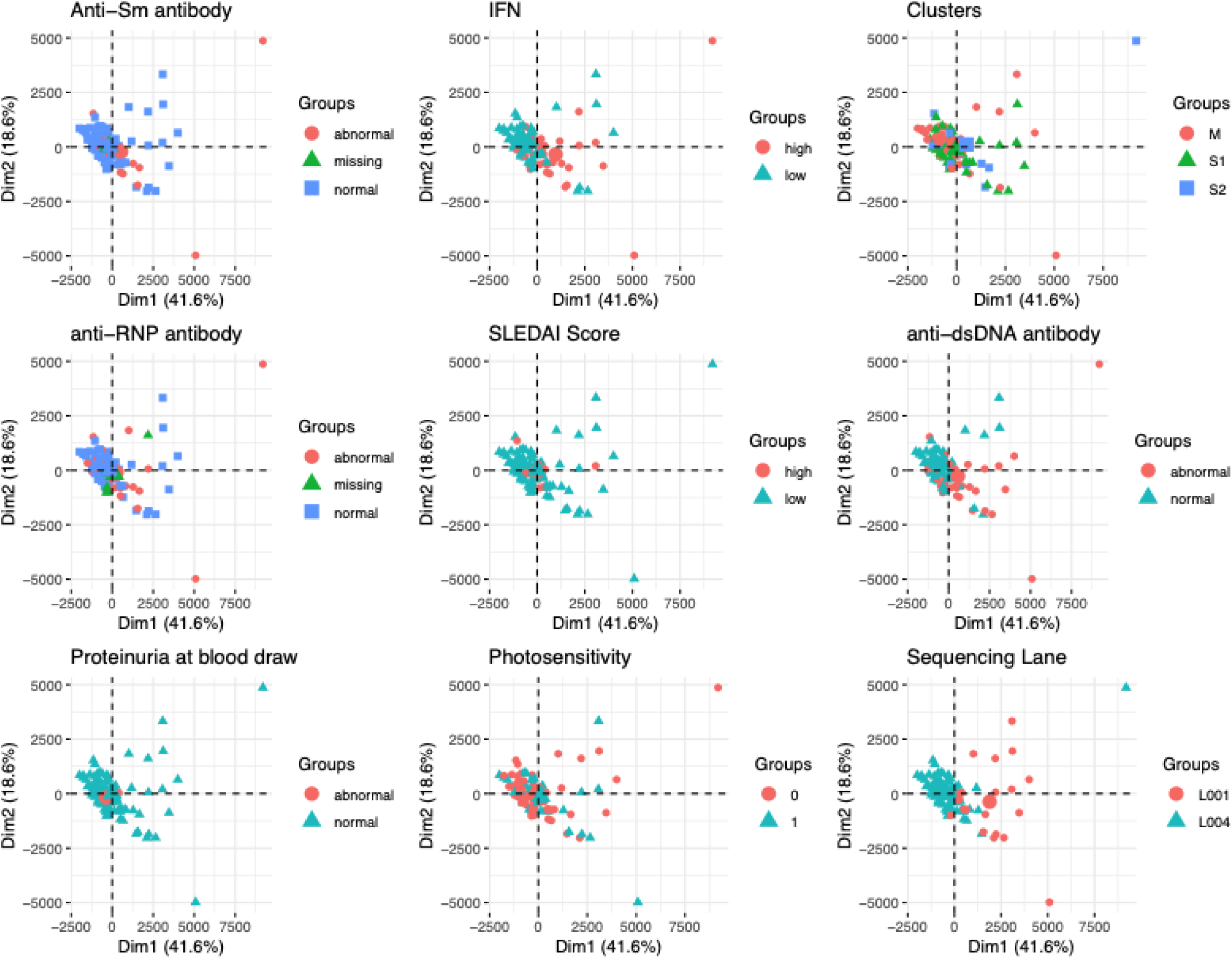
CD14 TE PCAs colored by demographics and phenotypes

**Supplementary Figure 4.**
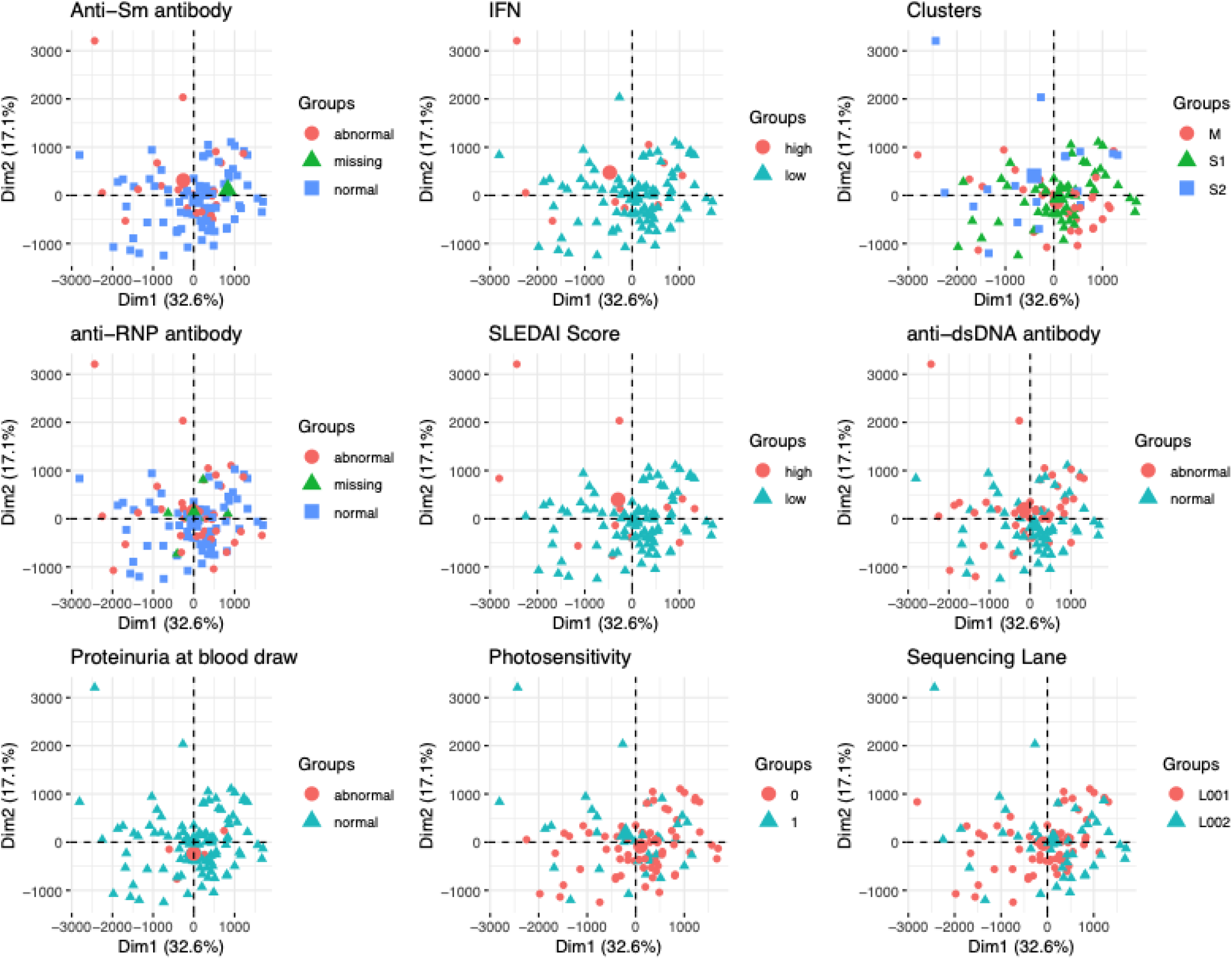
CD19 TE PCAs colored by demographics and phenotypes

**Supplementary Figure 5.**
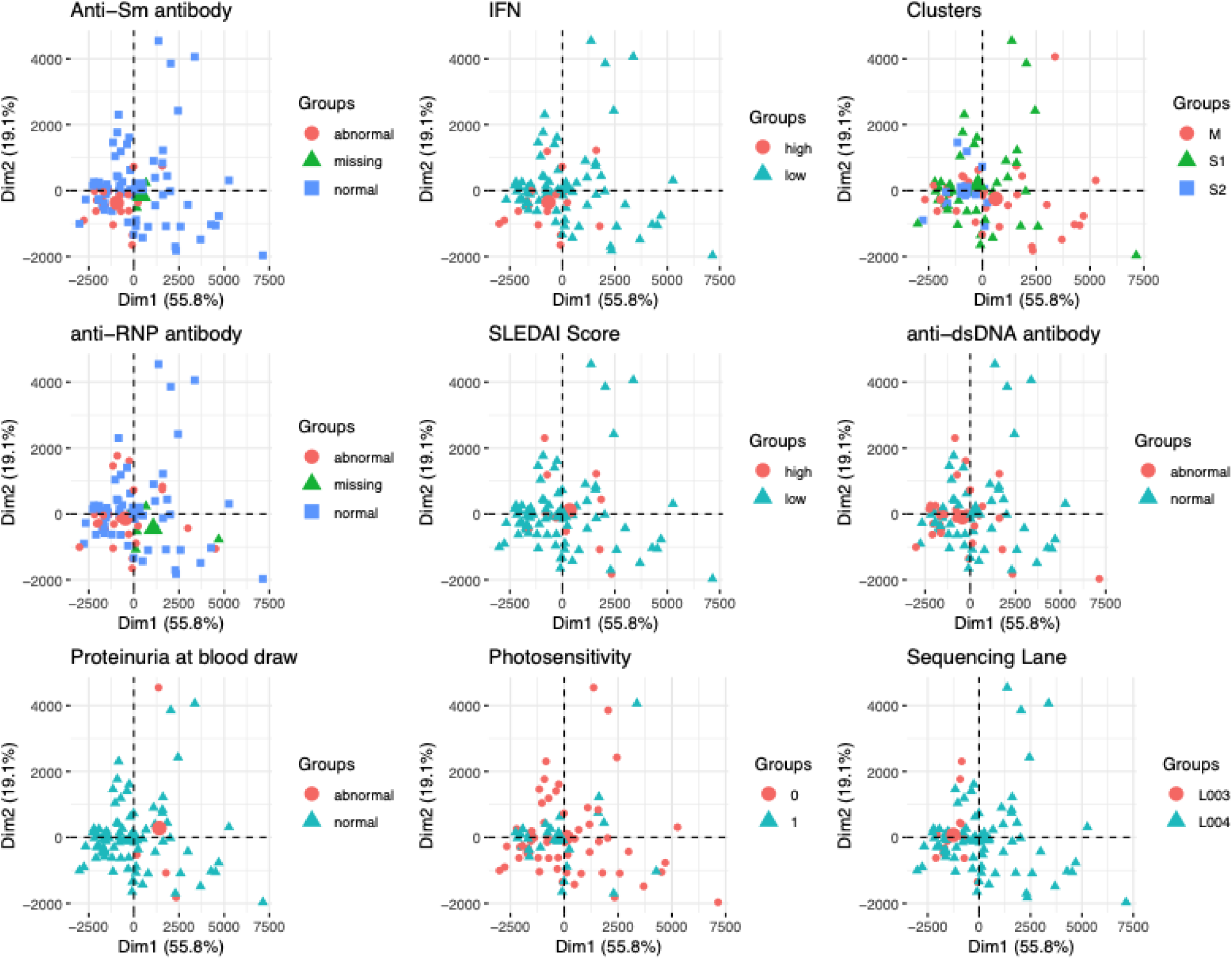
NK TE PCAs colored by demographics and SLE phenotypes

**Supplementary Figure 6.**
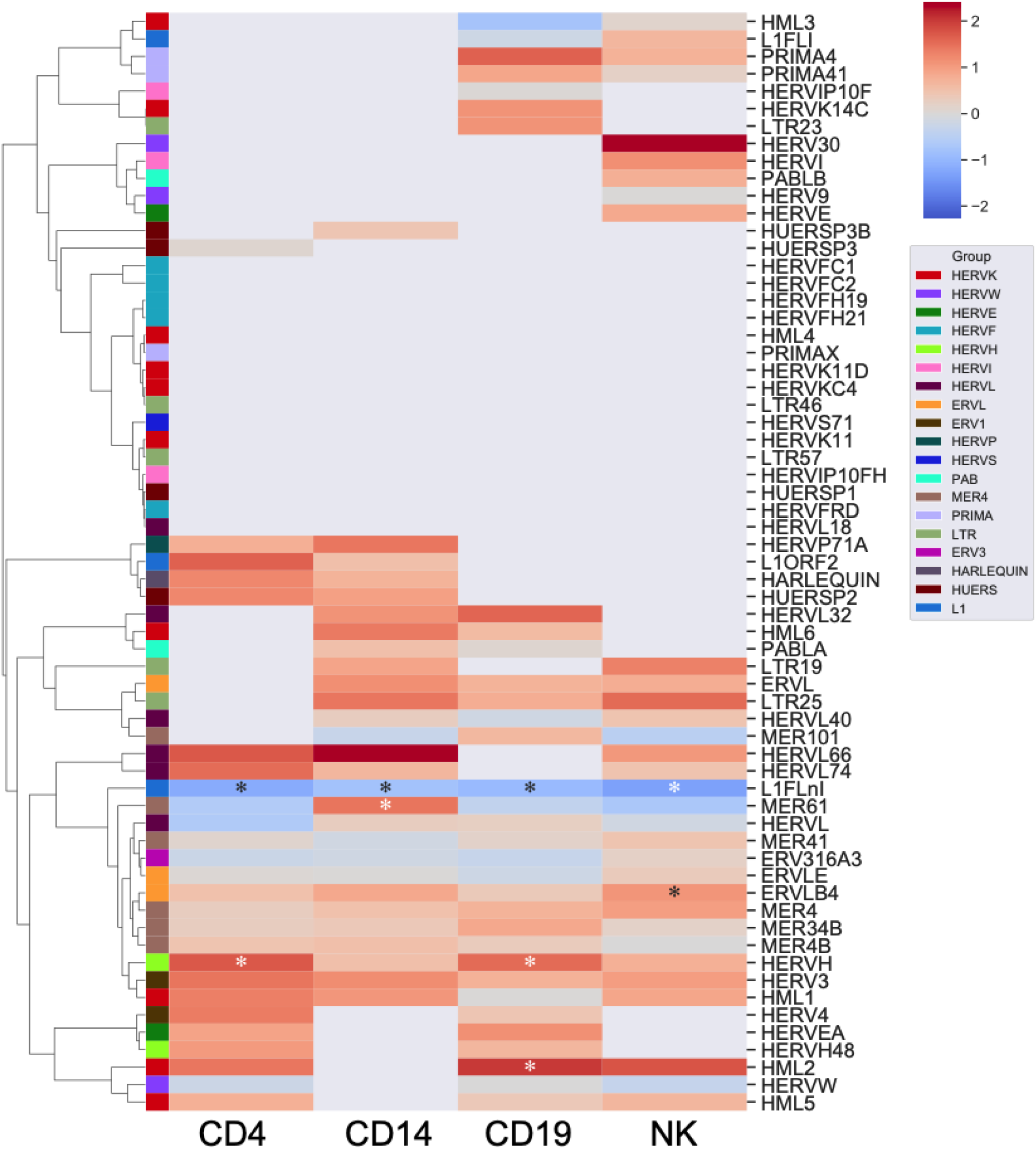
Family enrichment in combined TE DE analysis. Significant enrichment/depletion tested by hypergeometric test and marked with “*”.

**Supplementary Figure 7.**
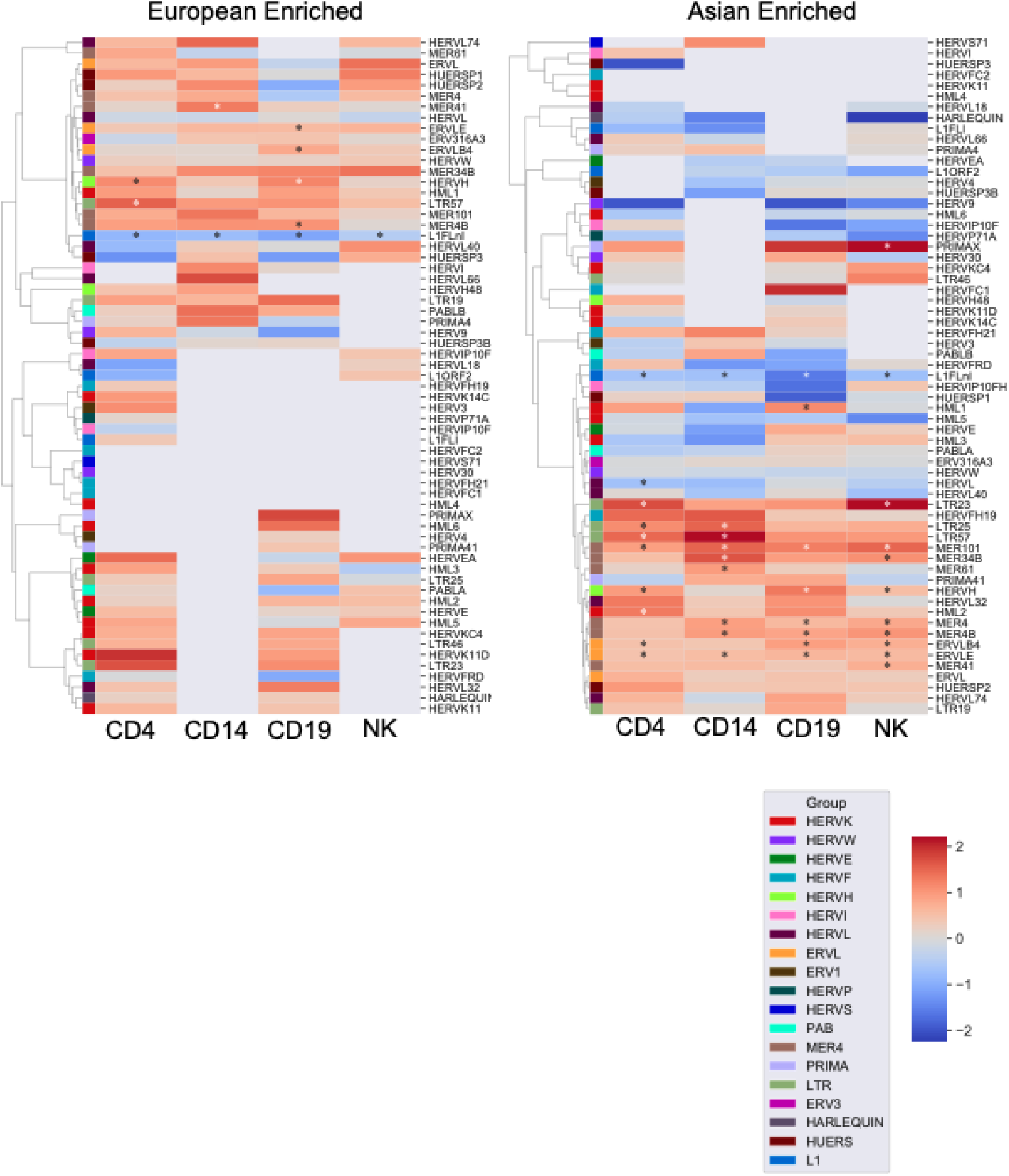
Family enrichment in ancestry stratified TE DE analysis. Significant enrichment/depletion tested by hypergeometric test and marked with “*”

**Supplementary Figure 8.**
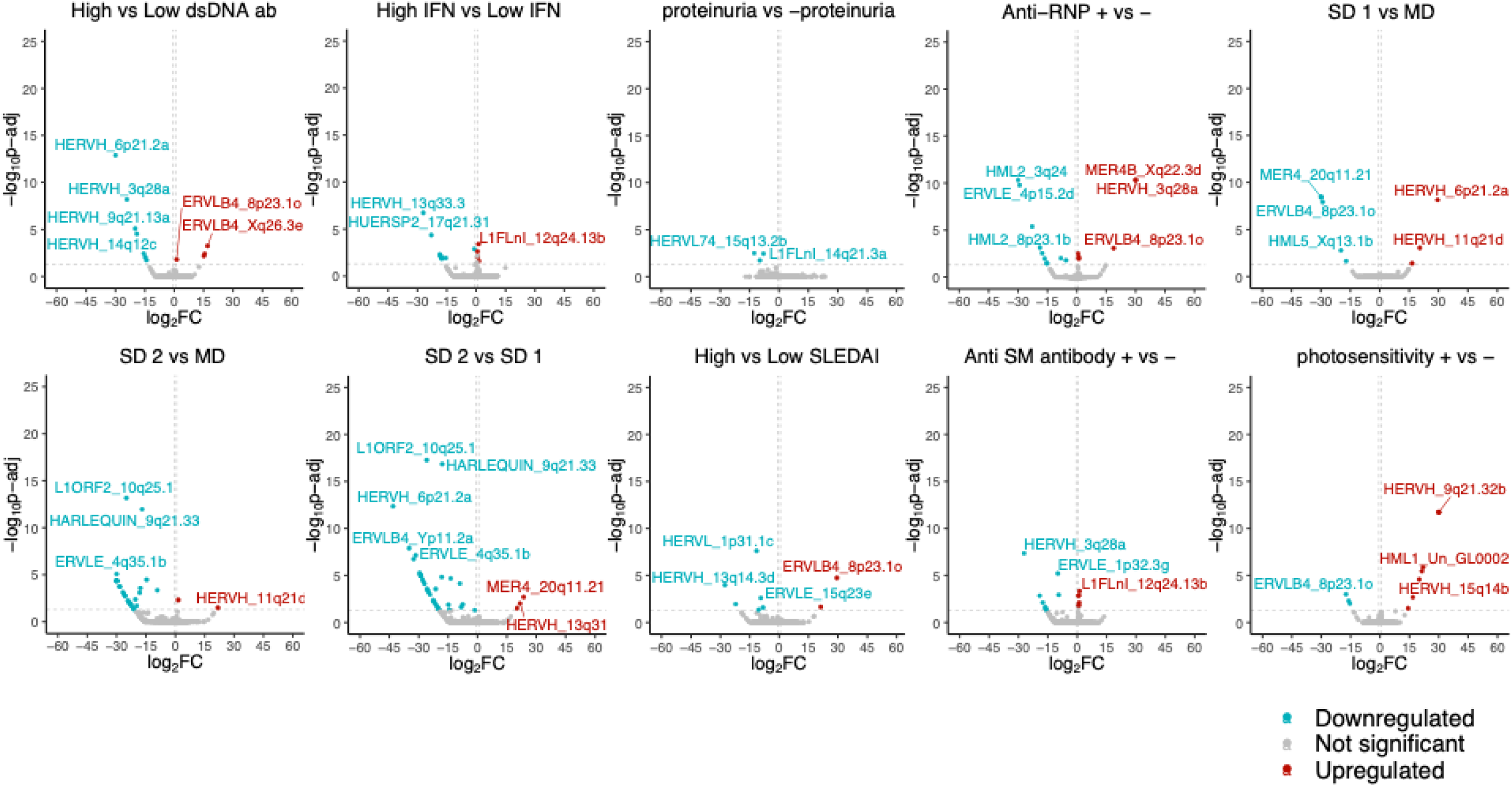
SLE subphenotype differential TE analysis for CD4 combined cohort

**Supplementary Figure 9.**
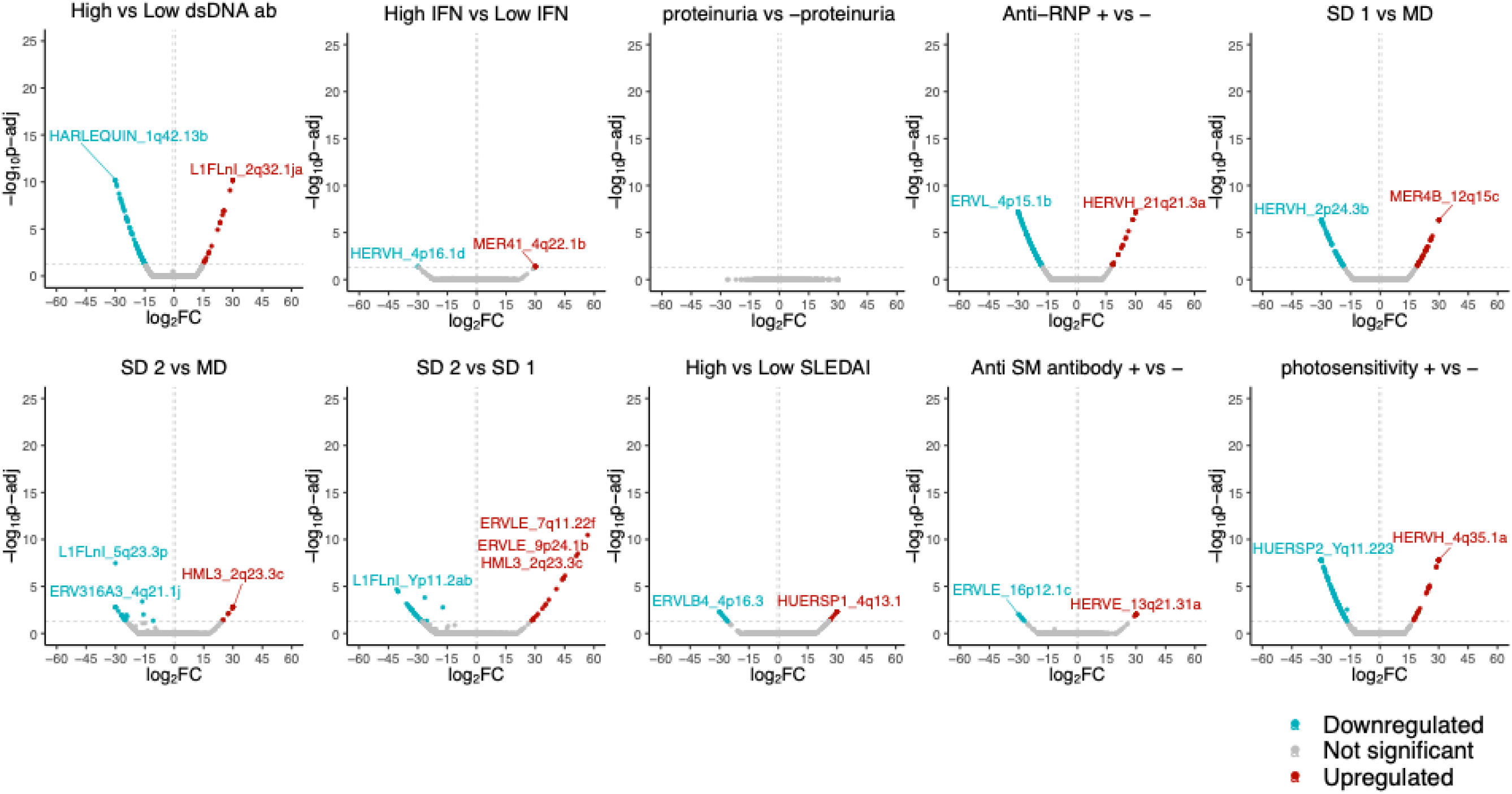
SLE subphenotype differential TE analysis for CD4 european enriched cohort

**Supplementary Figure 10.**
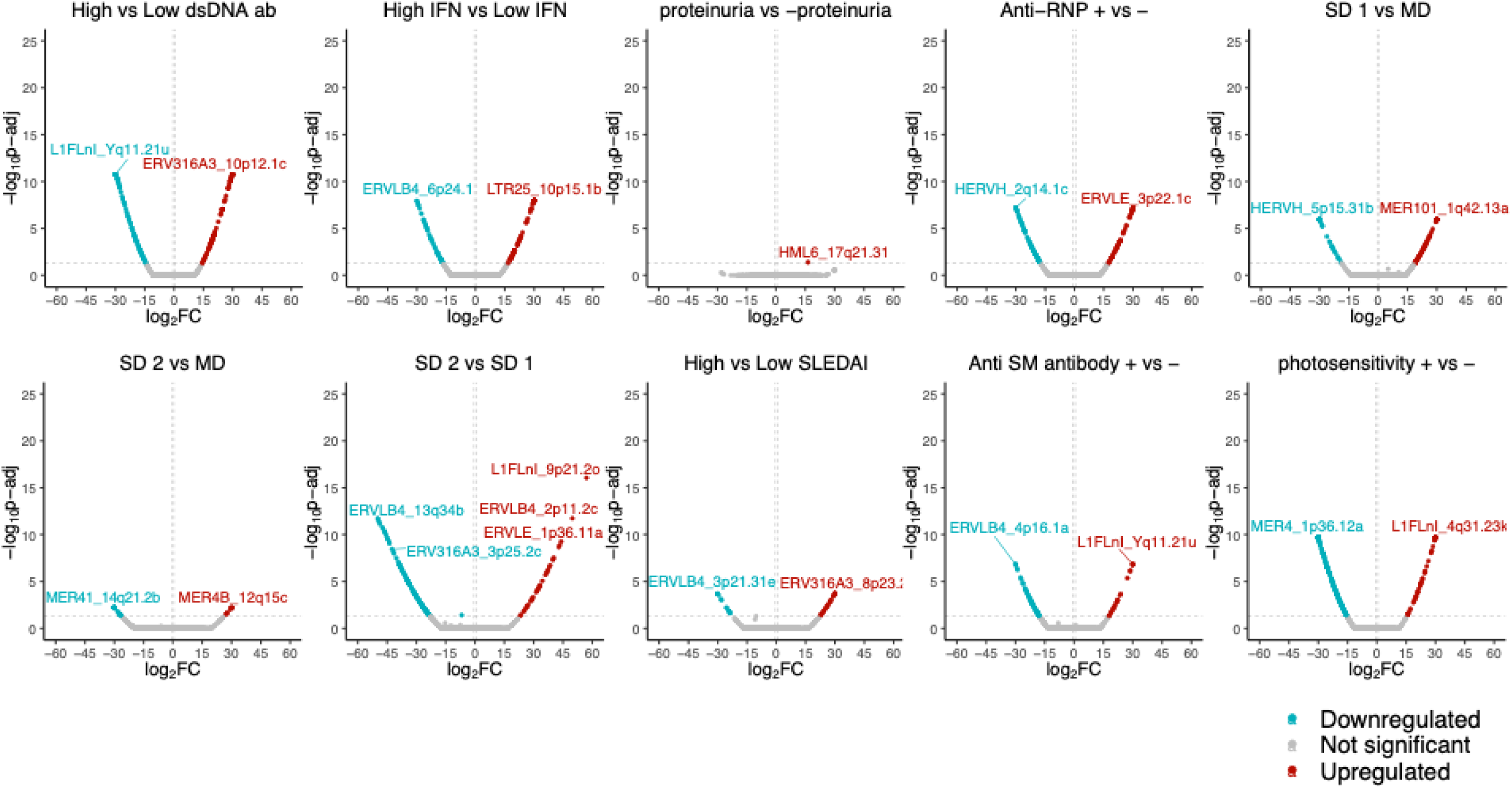
SLE subphenotype differential TE analysis for CD4 asian enriched cohort

**Supplementary Figure 11.**
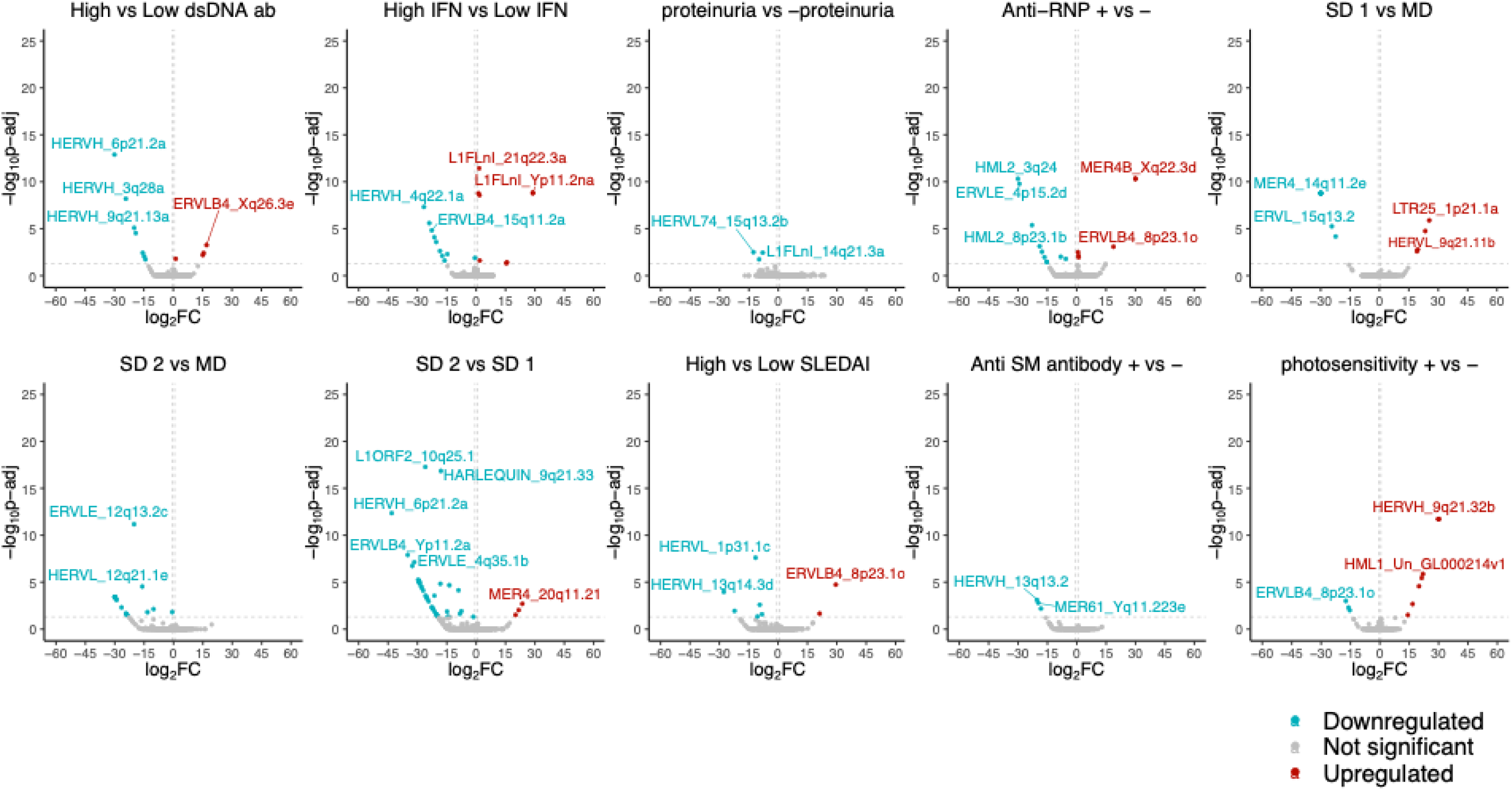
SLE subphenotype differential TE analysis for CD14 combined cohort

**Supplementary Figure 12.**
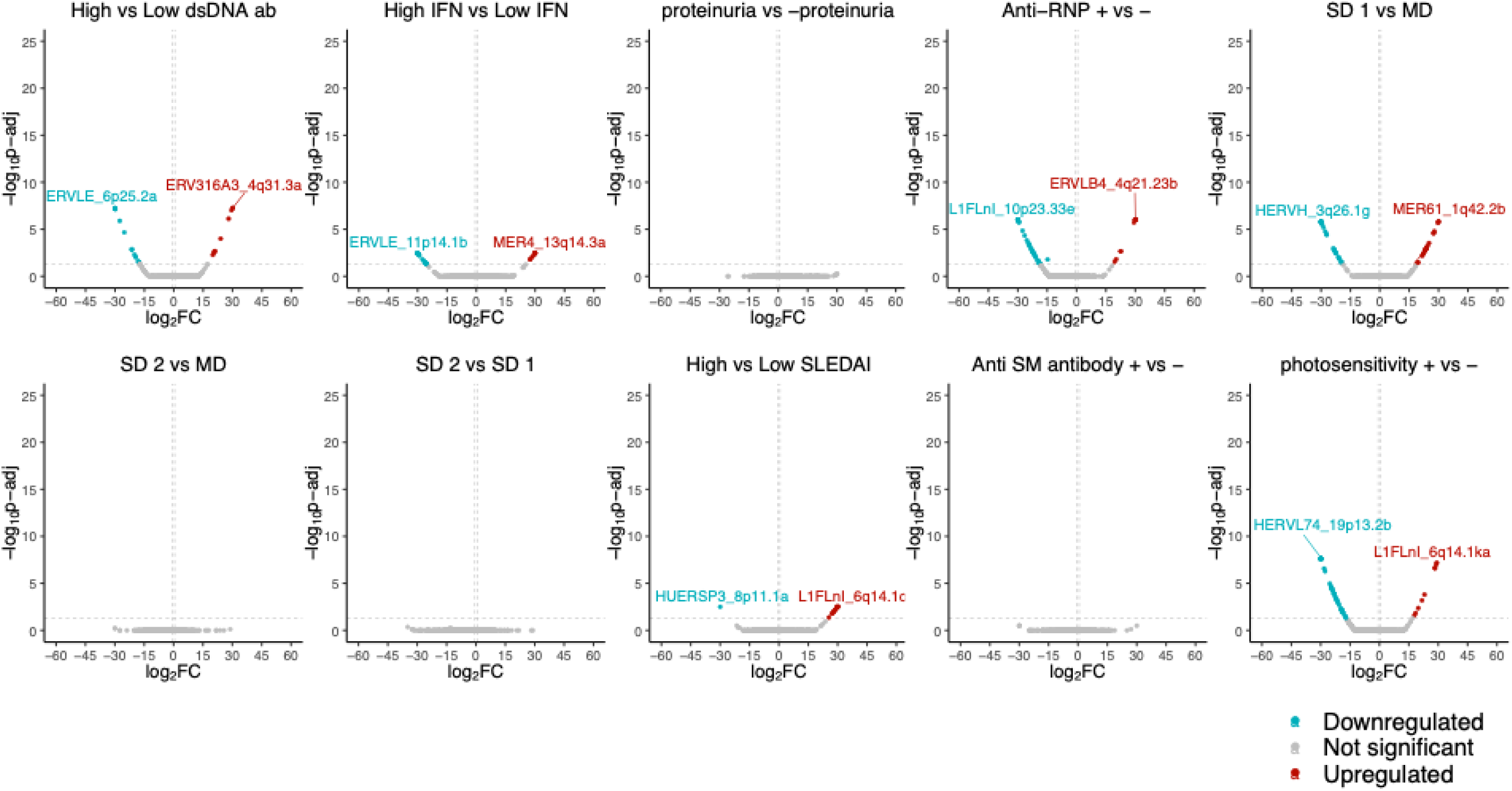
SLE subphenotype differential TE analysis for CD14 european enriched cohort

**Supplementary Figure 13.**
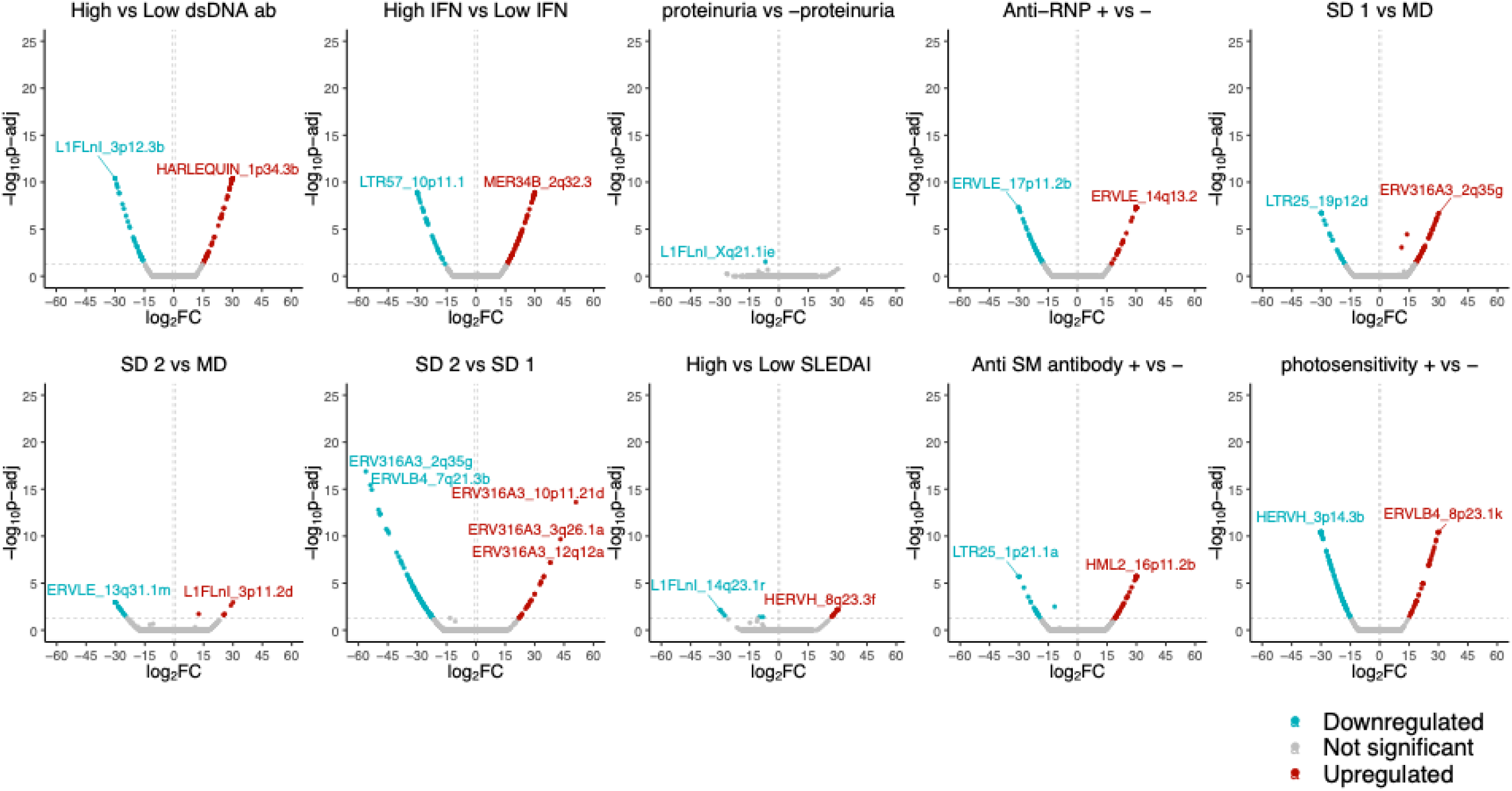
SLE subphenotype differential TE analysis for CD14 asian enriched cohort

**Supplementary Figure 14.**
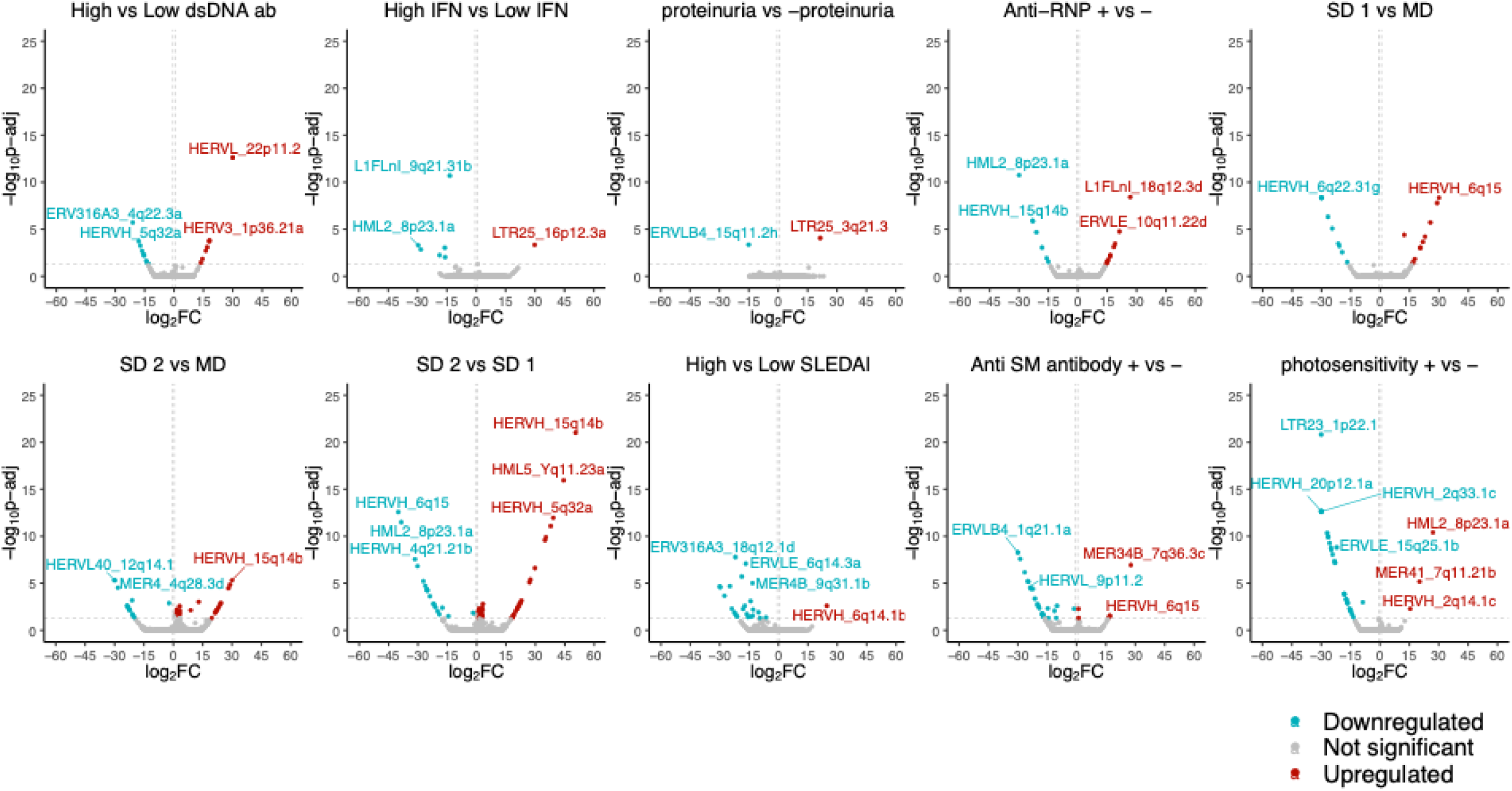
SLE subphenotype differential TE analysis for CD19 combined cohort

**Supplementary Figure 15.**
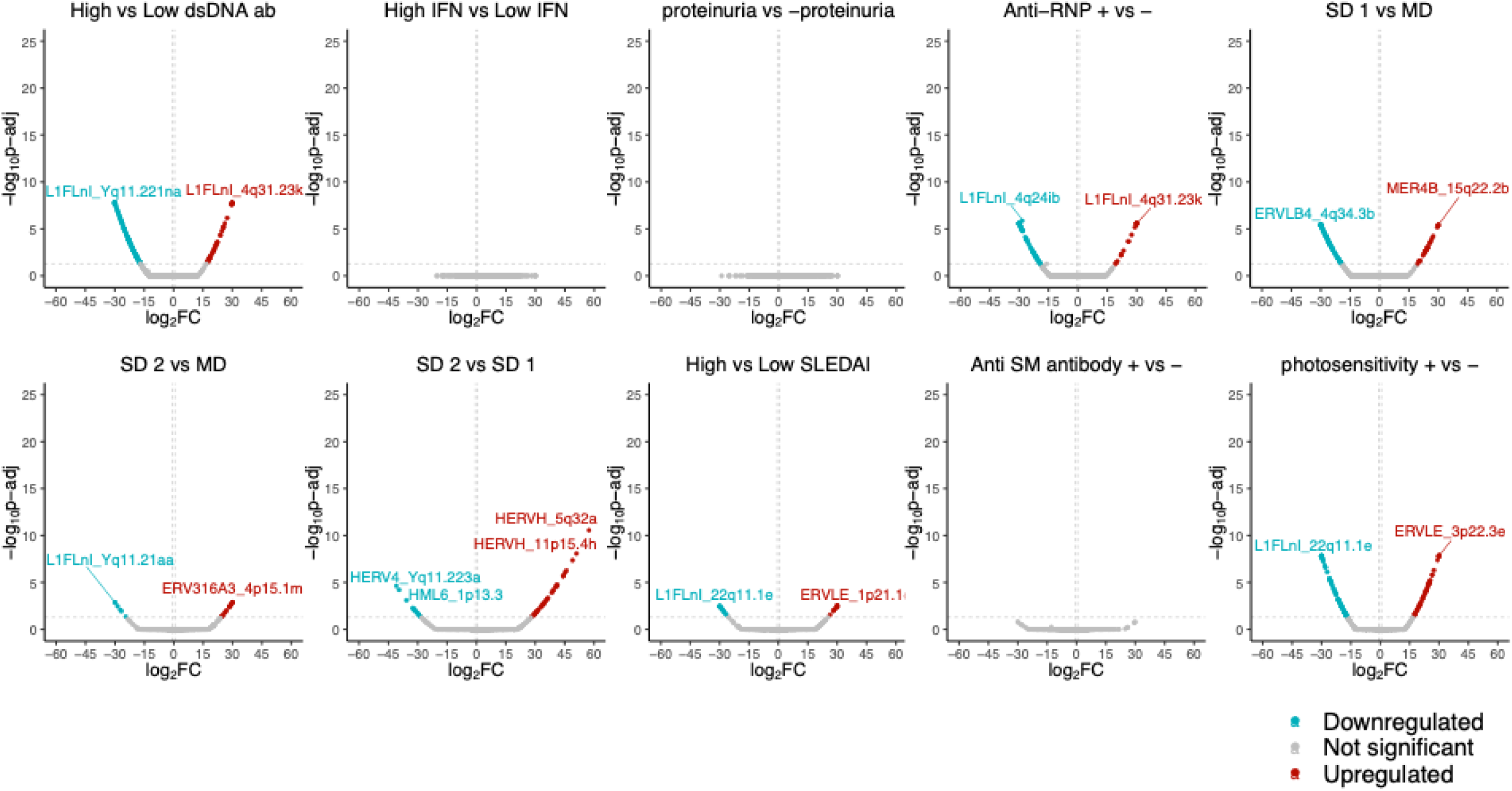
SLE subphenotype differential TE analysis for CD19 european enriched cohort

**Supplementary Figure 16.**
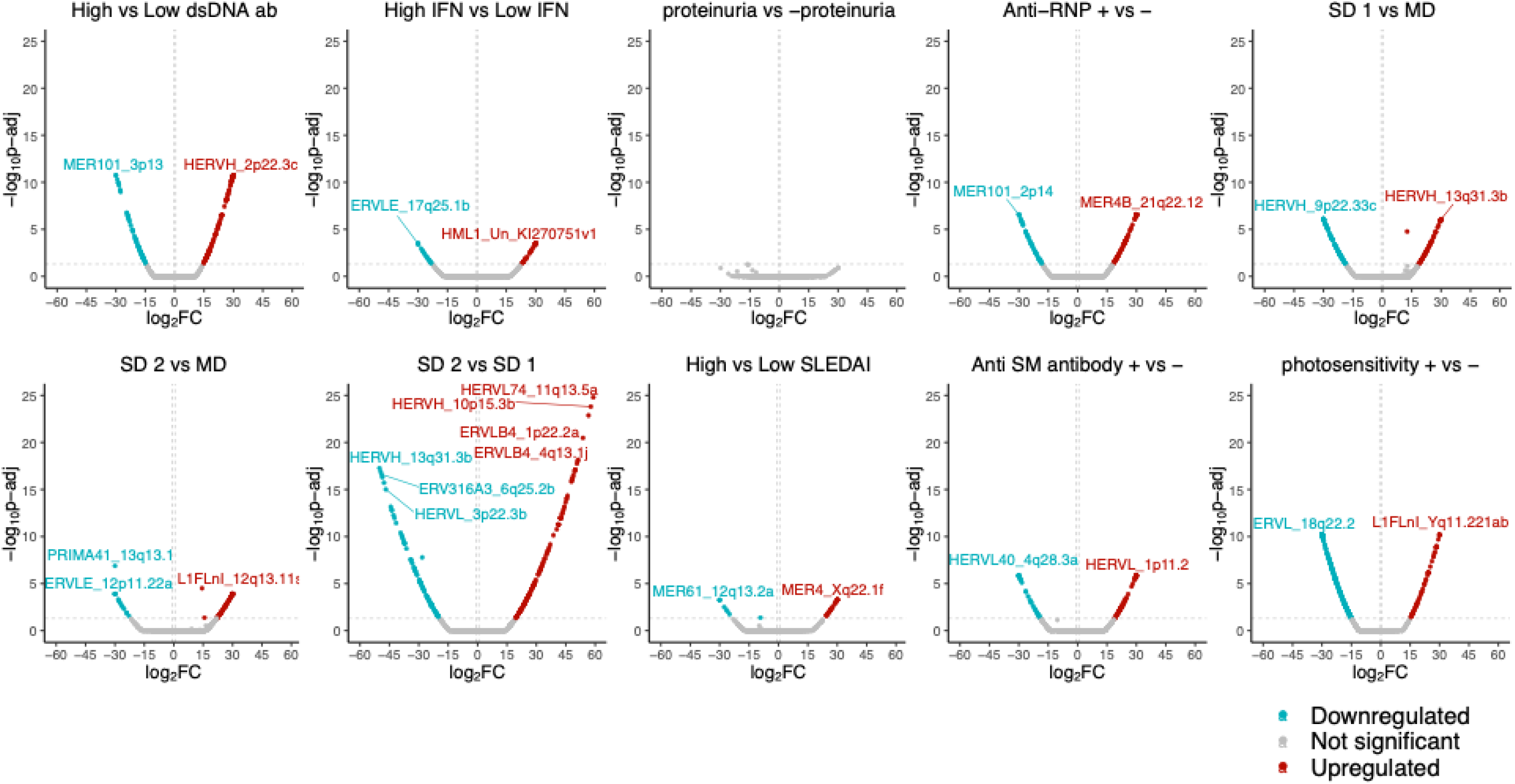
SLE subphenotype differential TE analysis for CD19 asian enriched cohort

**Supplementary Figure 17.**
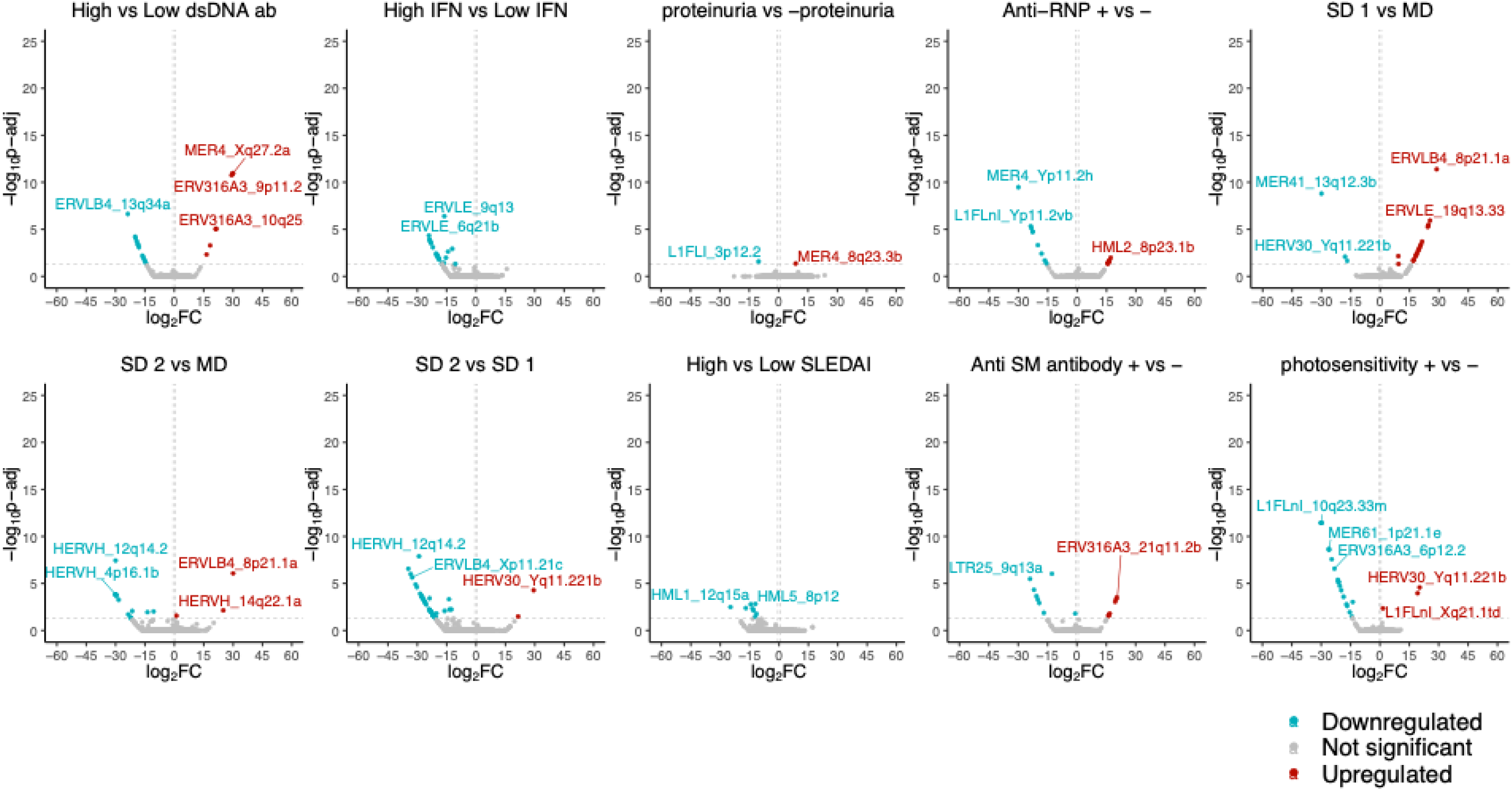
SLE subphenotype differential TE analysis for NK combined cohort

**Supplementary Figure 18.**
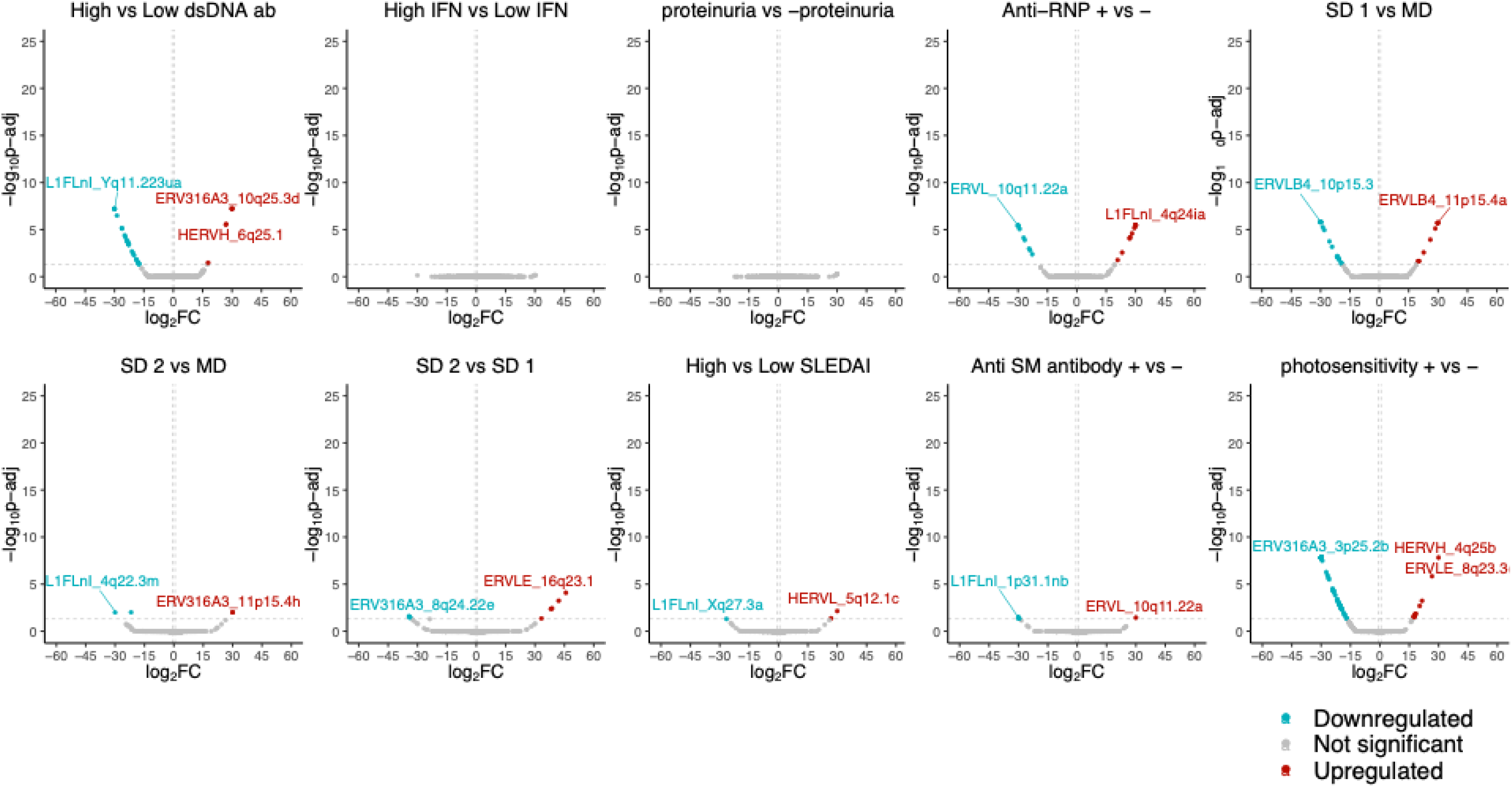
SLE subphenotype differential TE analysis for NK european enriched cohort

**Supplementary Figure 19.**
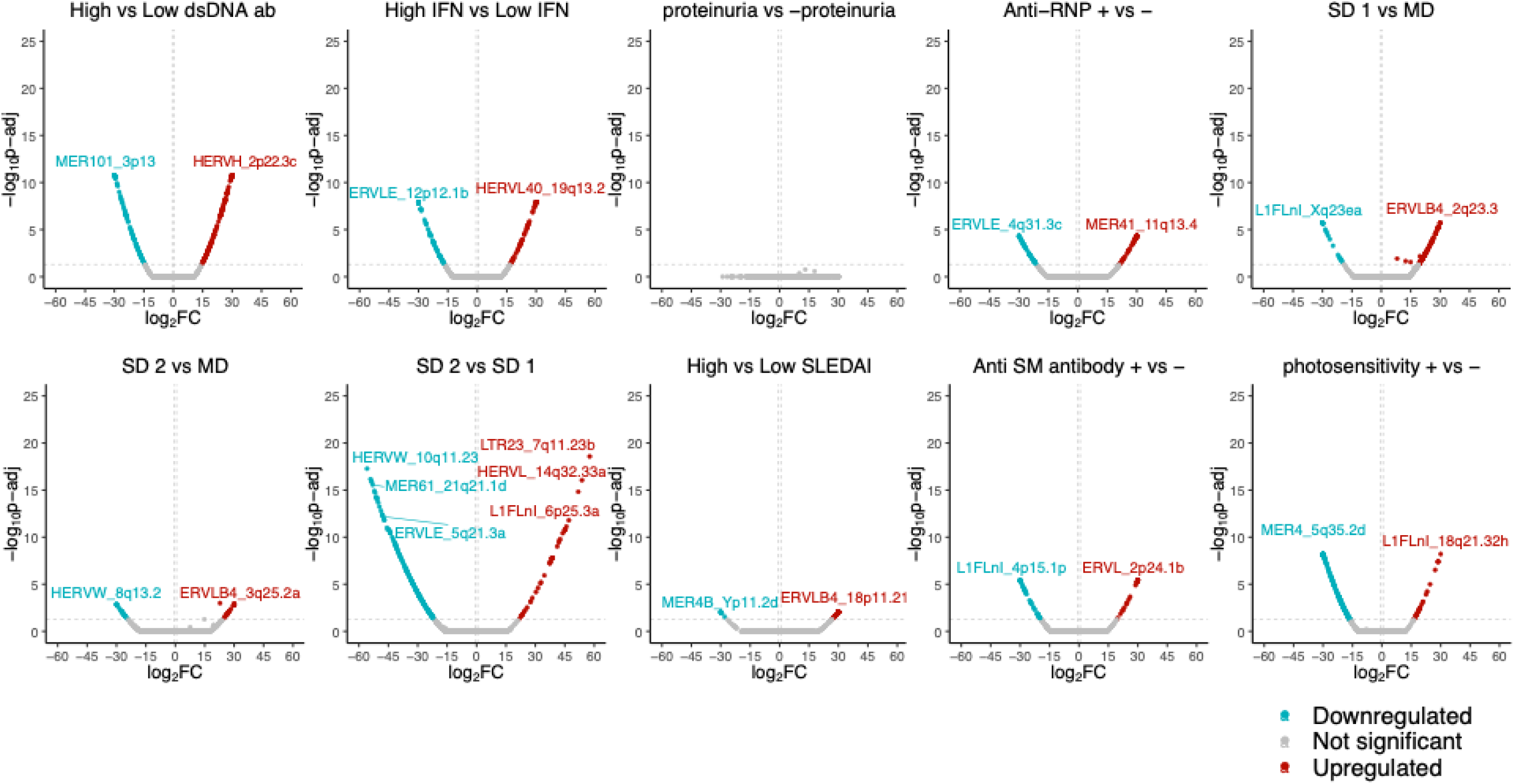
SLE subphenotype differential TE analysis for NK asian enriched cohort

