## Supplementary Figures for "Cell-Specific Transposable Element Gene Expression Analysis Identifies Associations with Systemic Lupus Erythematosus Phenotypes"

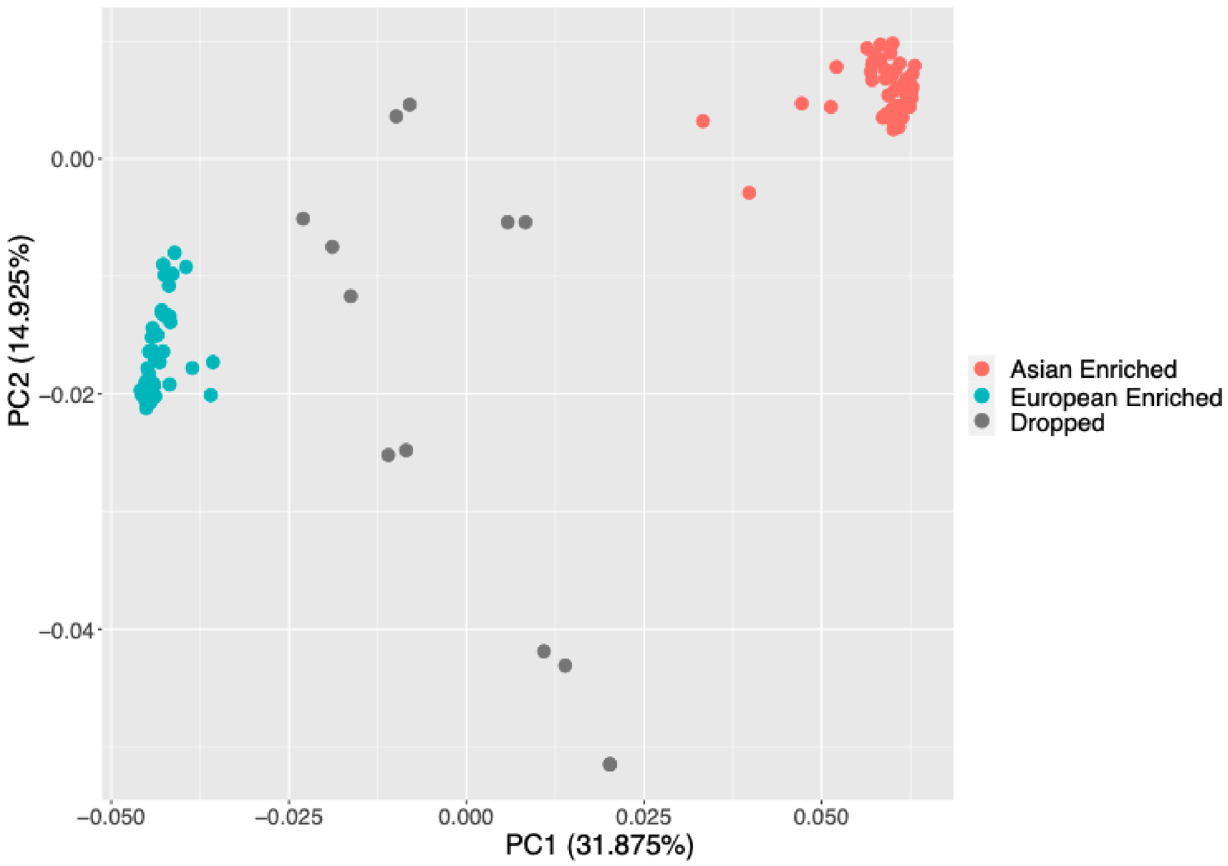


**Supplementary Figure 1.** PCA plot of genotyping data across 120 patients. Individuals are colored by ancestry. Individuals in gray are considered admixed and were dropped from the stratified analysis.

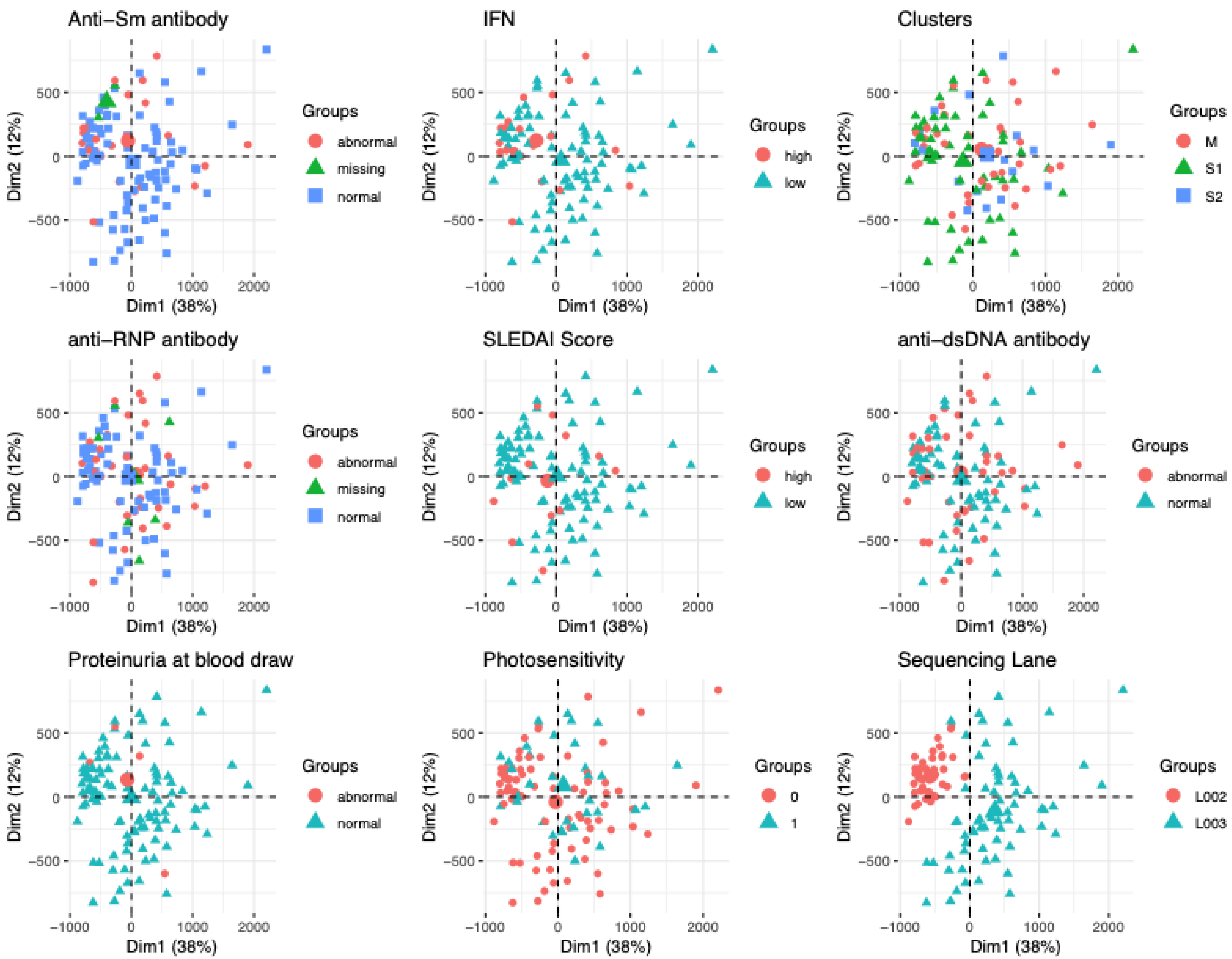


**Supplementary Figure 2.** CD4 TE PCAs colored by demographics and phenotypes

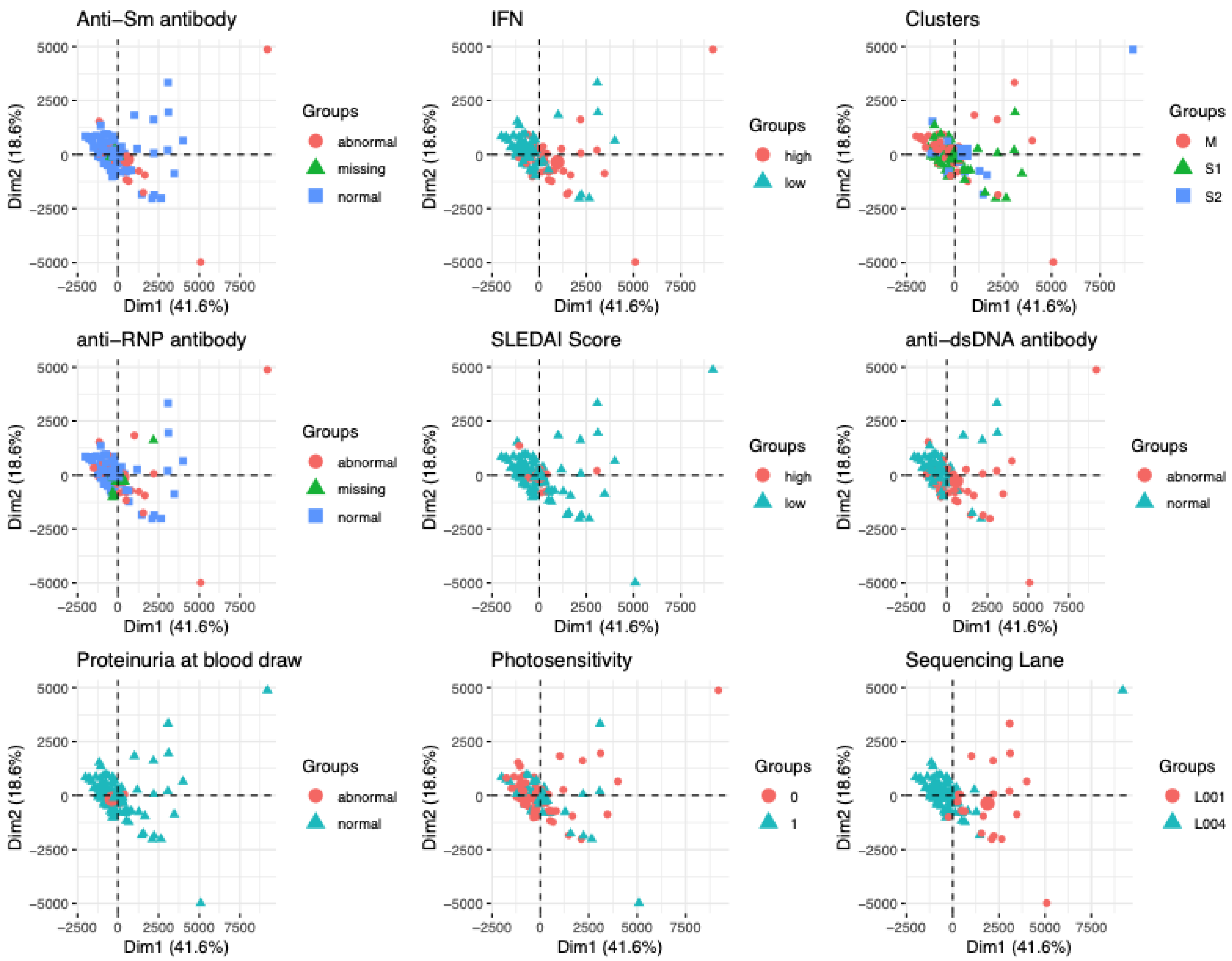


**Supplementary Figure 3.** CD14 TE PCAs colored by demographics and phenotypes

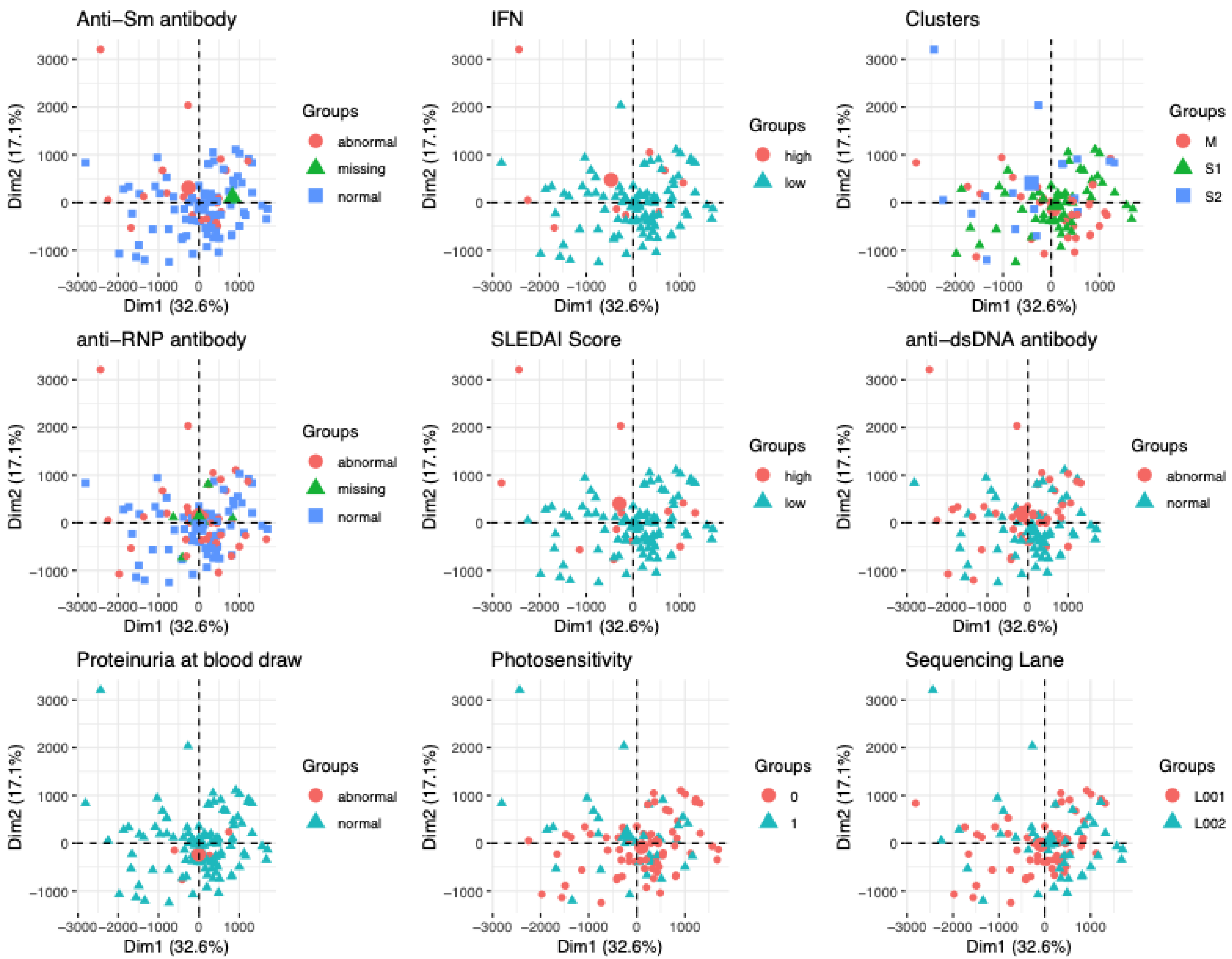


**Supplementary Figure 4.** CD19 TE PCAs colored by demographics and phenotypes

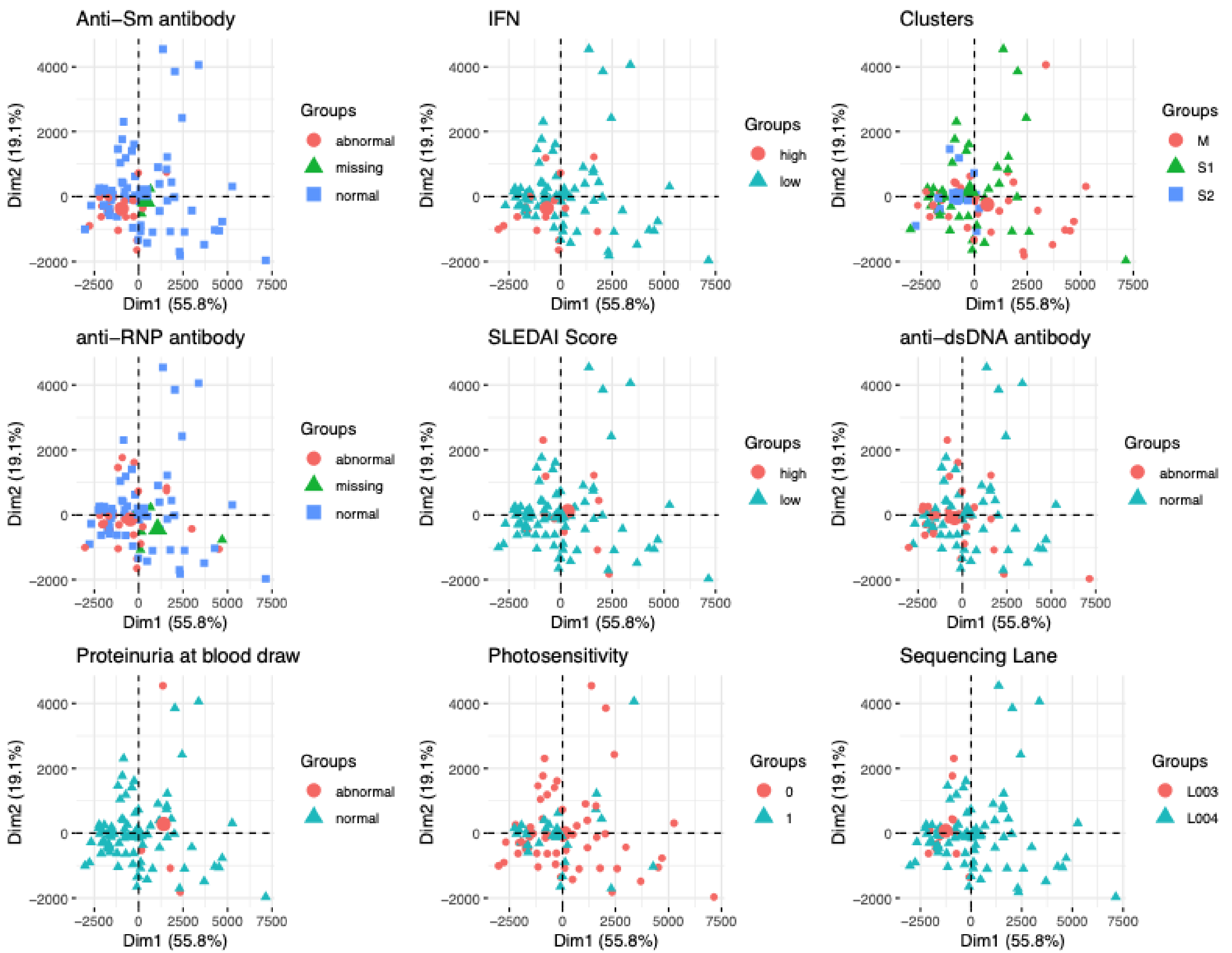


**Supplementary Figure 5.** NK TE PCAs colored by demographics and SLE phenotypes

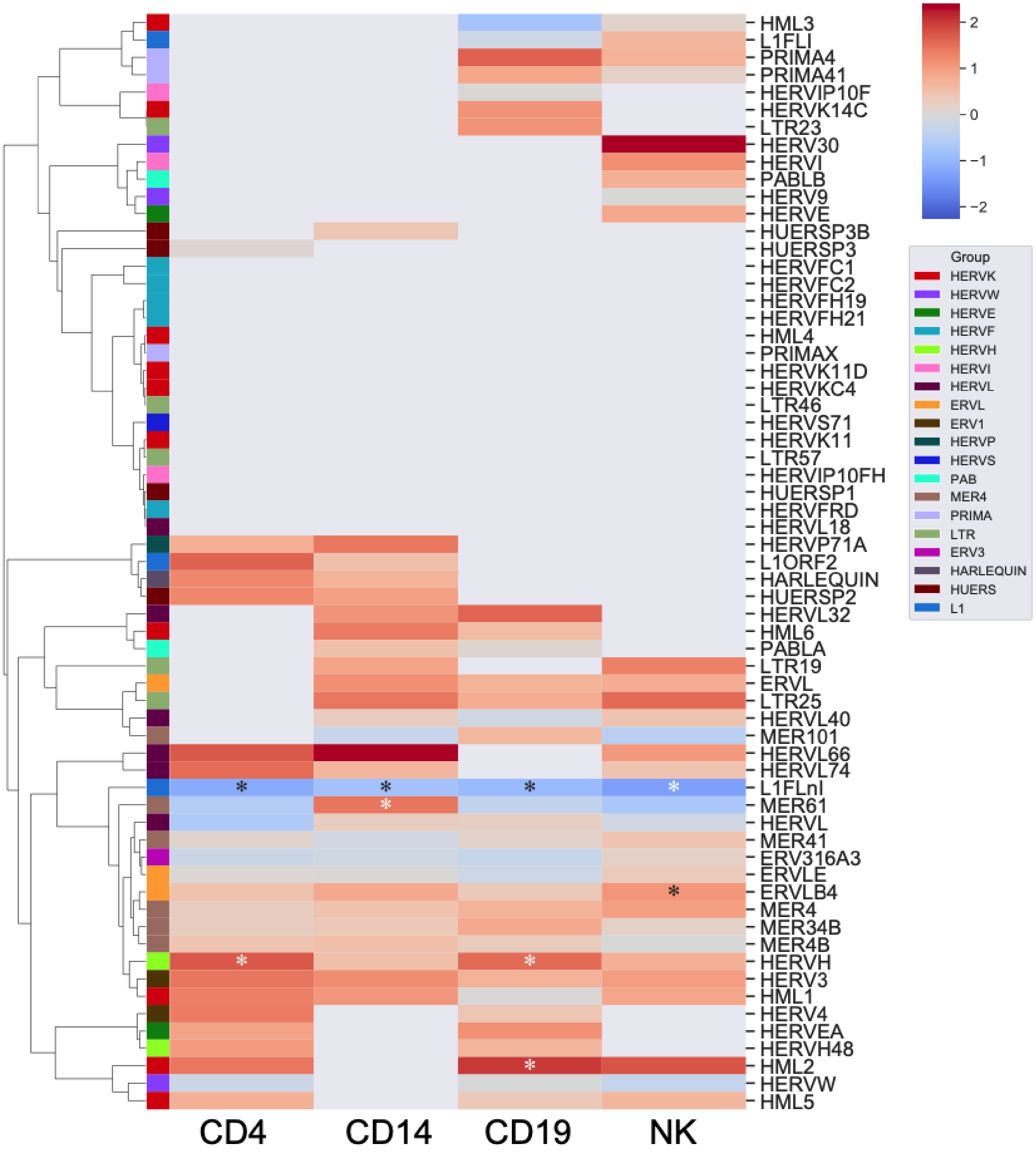


**Supplementary Figure 6.** Family enrichment in combined TE DE analysis. Significant enrichment/depletion tested by hypergeometric test and marked with "*".

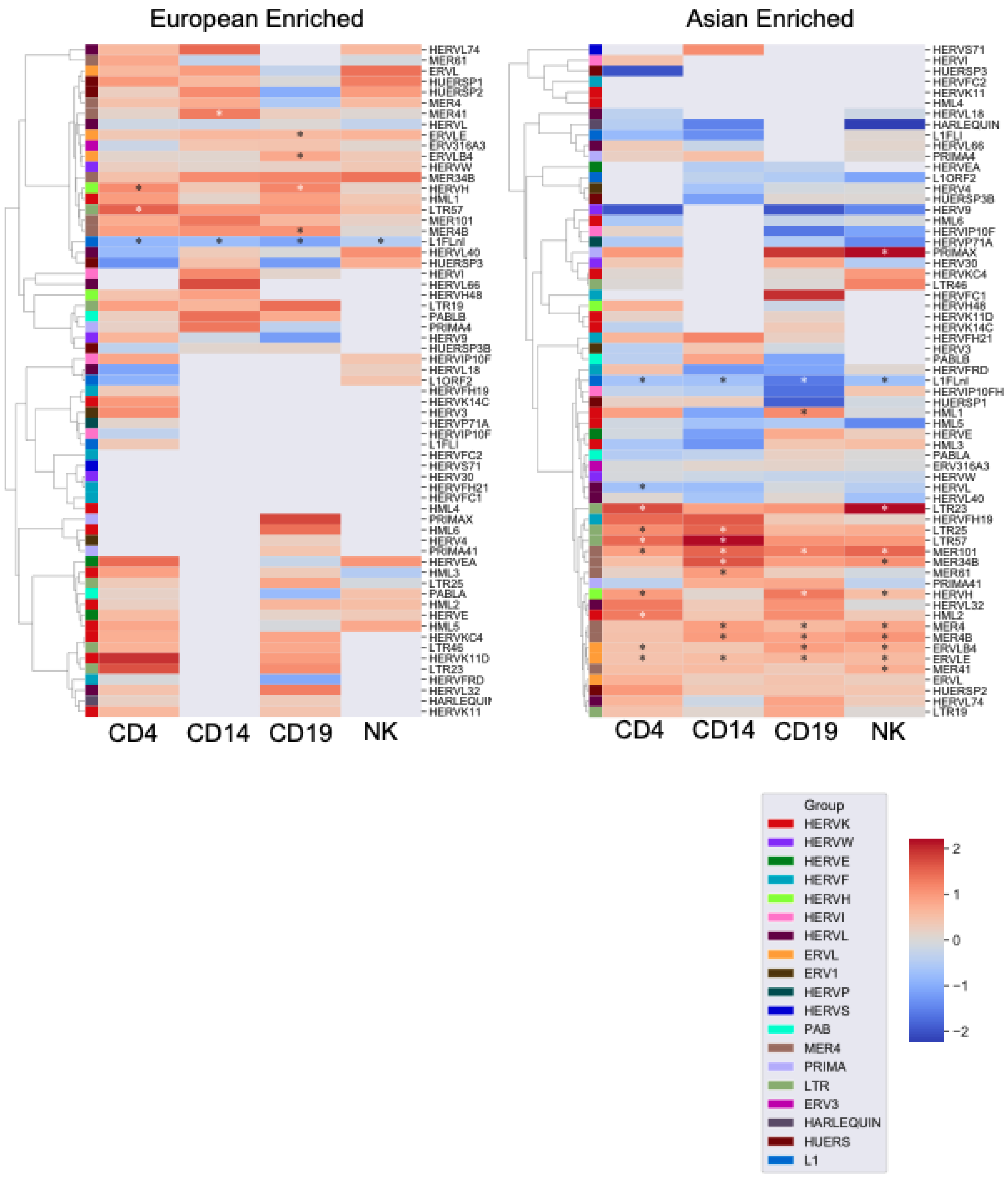


**Supplementary Figure 7.** Family enrichment in ancestry stratified TE DE analysis. Significant enrichment/depletion tested by hypergeometric test and marked with “*”

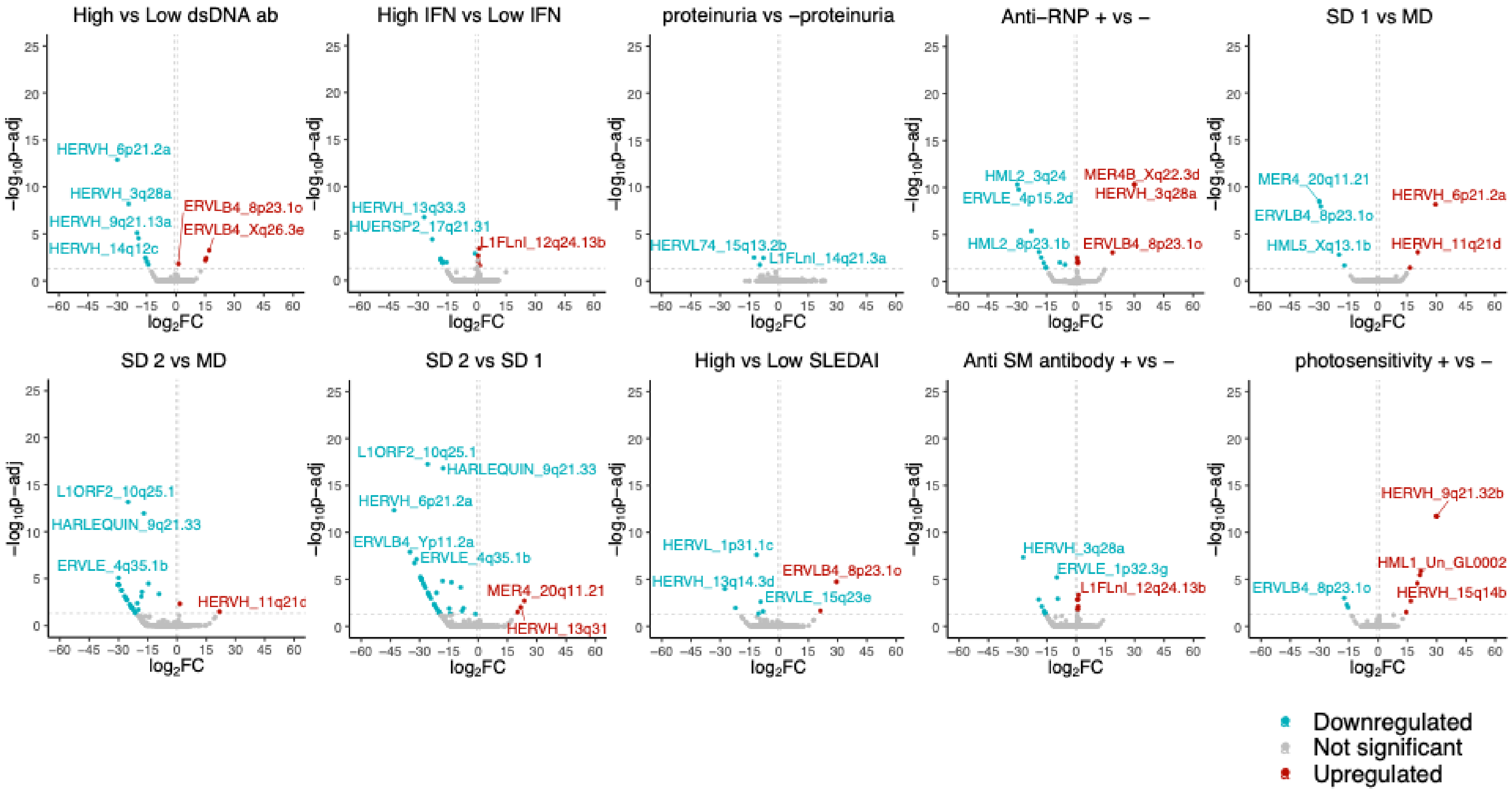


**Supplementary Figure 8.** SLE subphenotype differential TE analysis for CD4 combined cohort

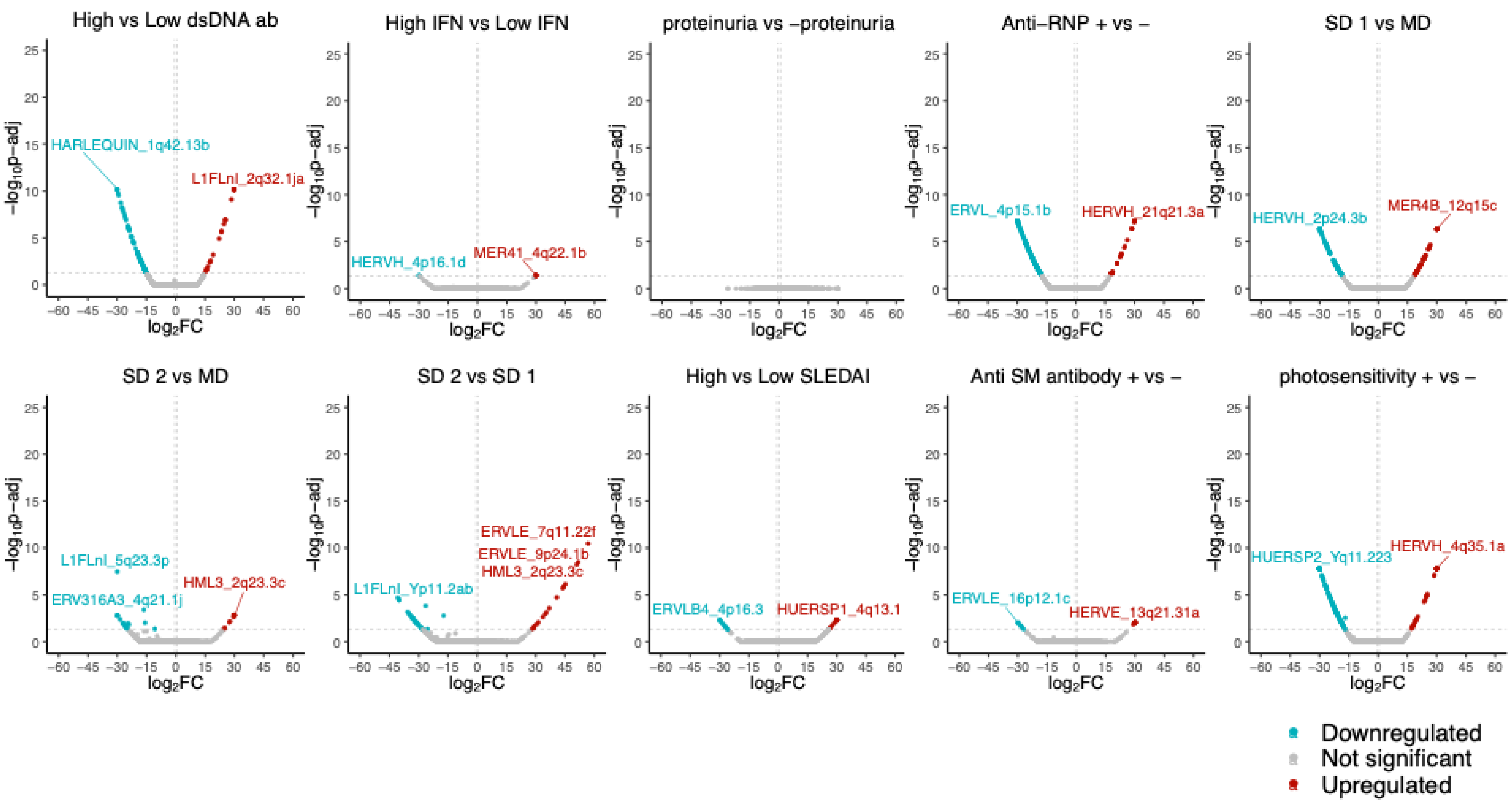


**Supplementary Figure 9.** SLE subphenotype differential TE analysis for CD4 european enriched cohort

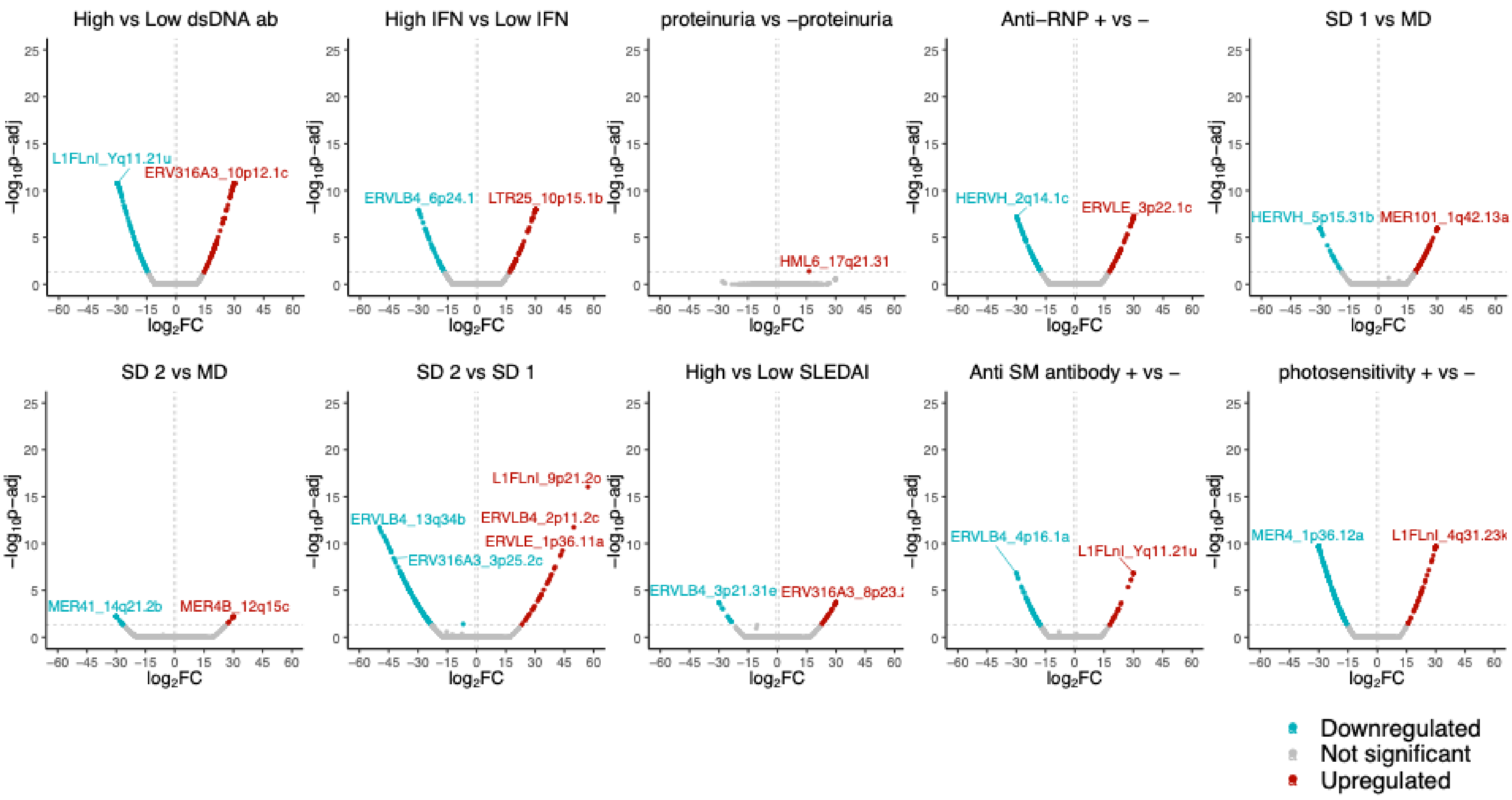


**Supplementary Figure 10.** SLE subphenotype differential TE analysis for CD4 asian enriched cohort

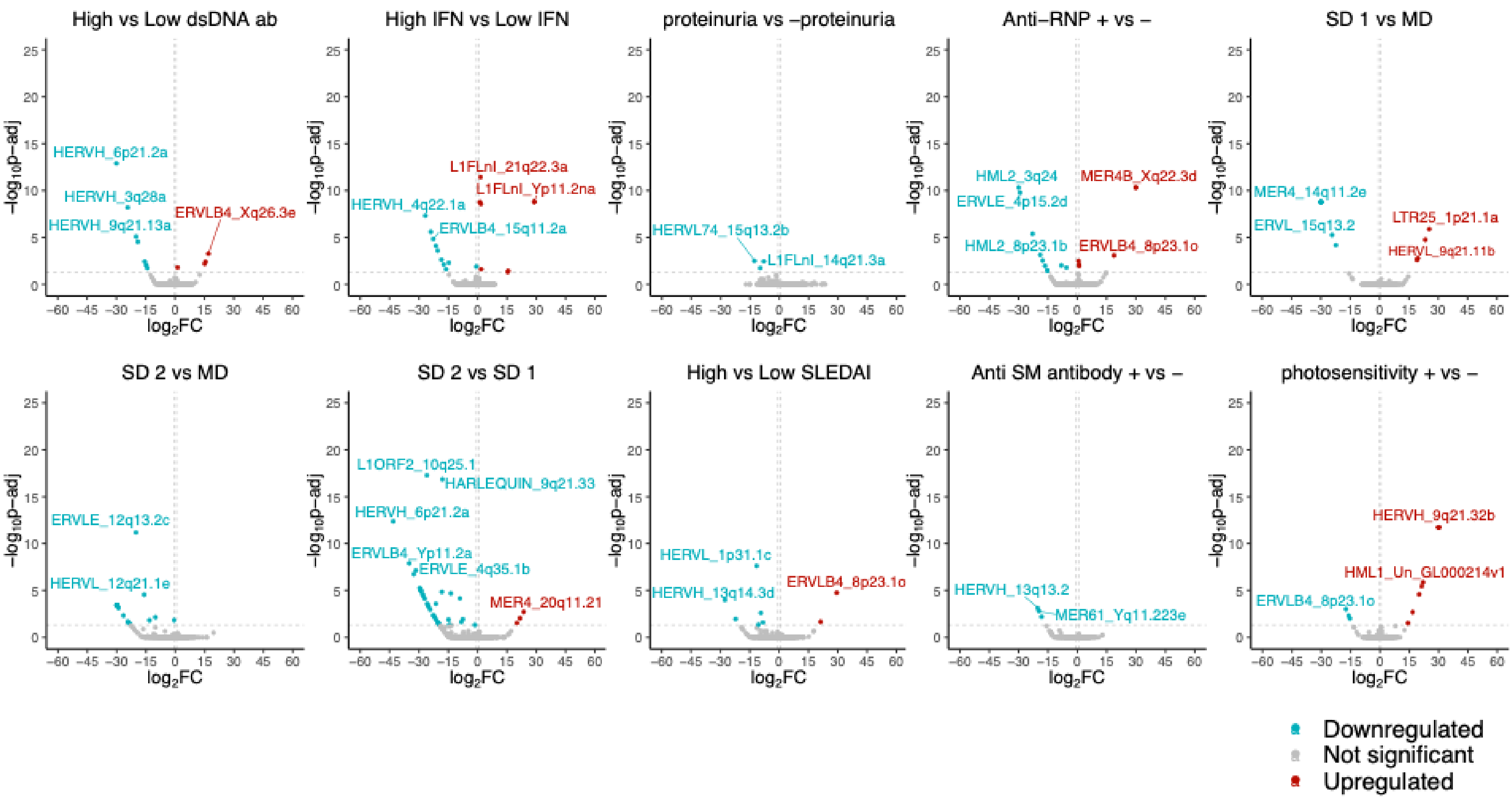


**Supplementary Figure 11.** SLE subphenotype differential TE analysis for CD14 combined cohort

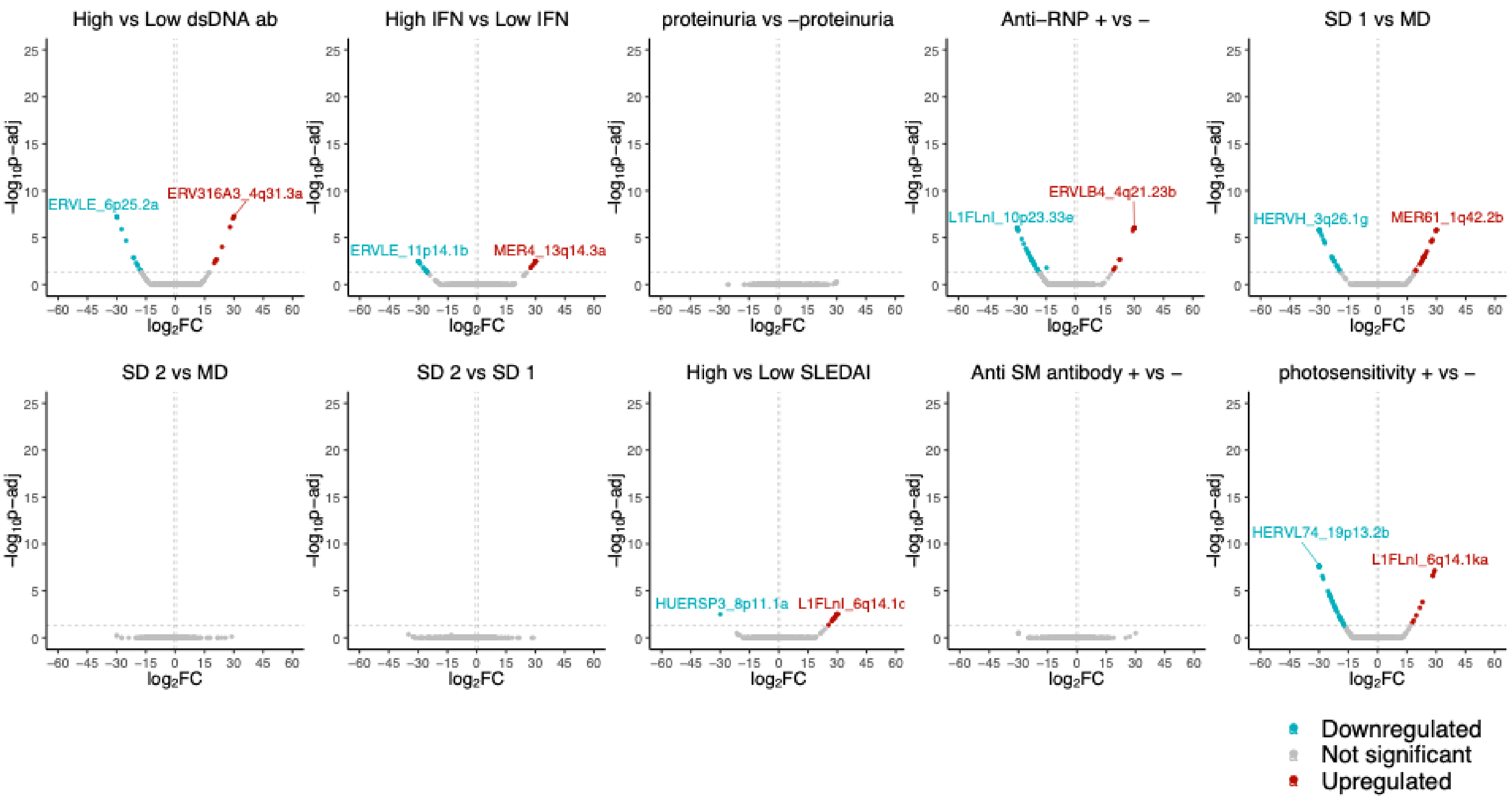


**Supplementary Figure 12.** SLE subphenotype differential TE analysis for CD14 european enriched cohort

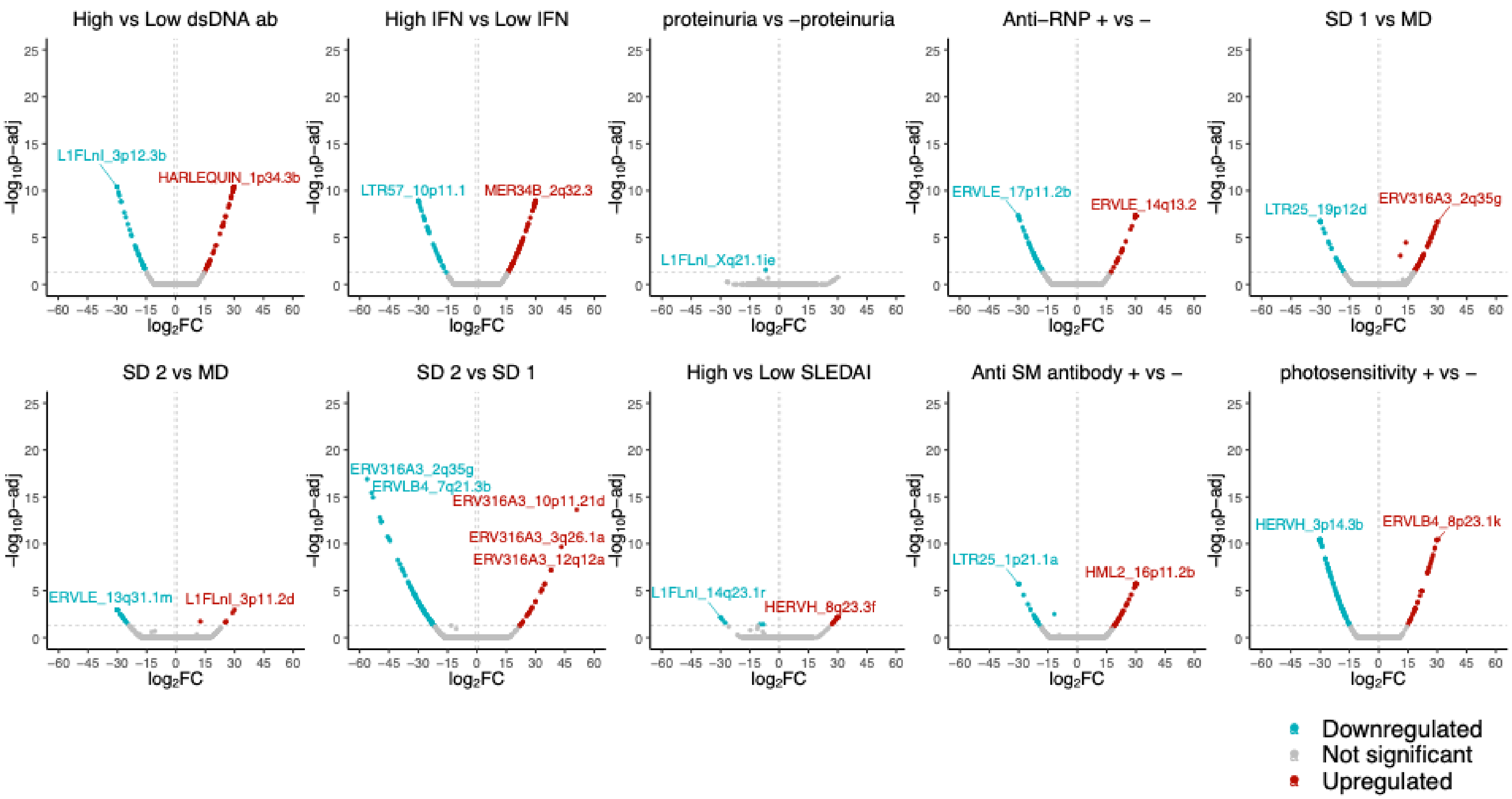


**Supplementary Figure 13.** SLE subphenotype differential TE analysis for CD14 asian enriched cohort

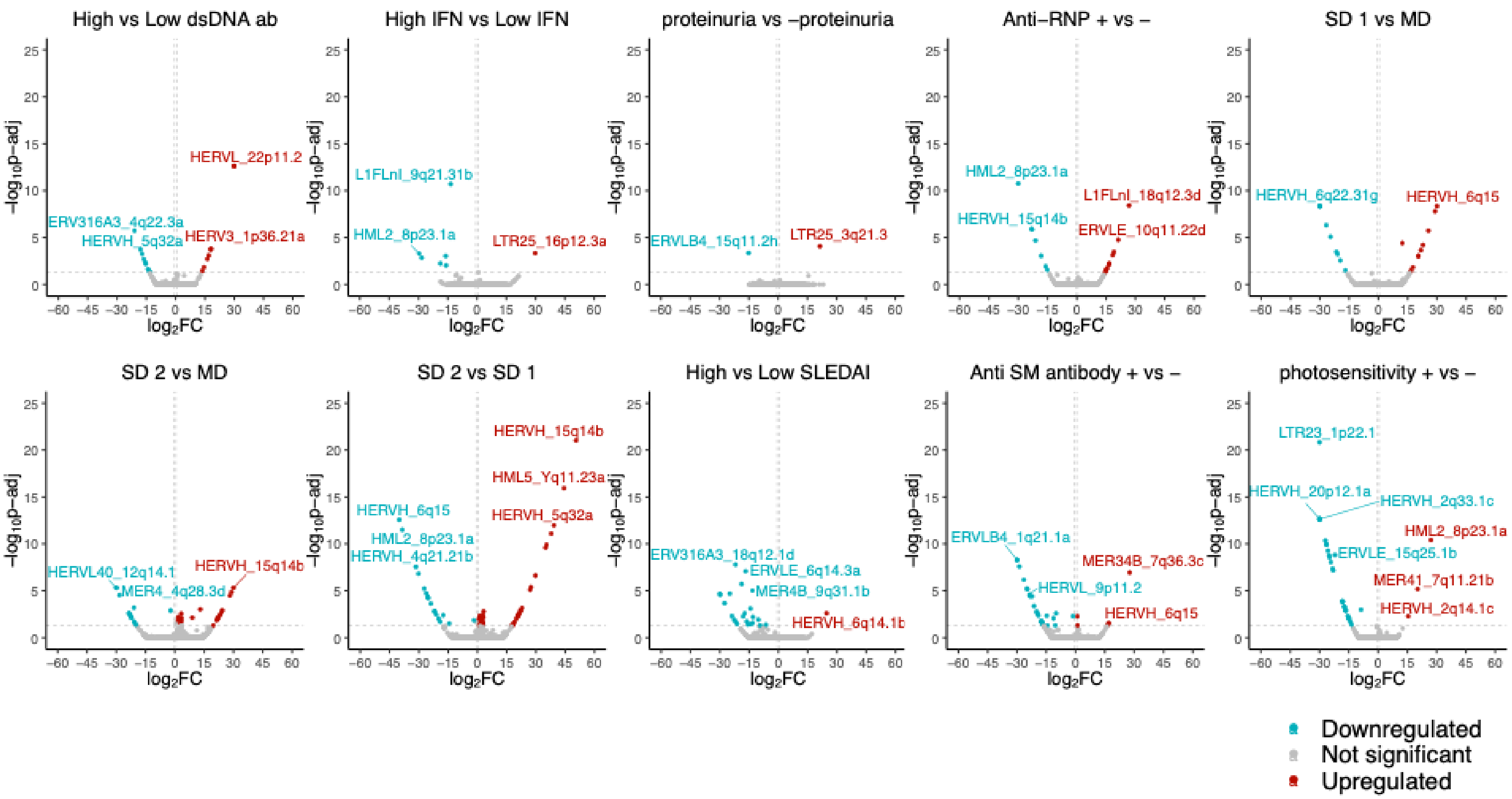


**Supplementary Figure 14.** SLE subphenotype differential TE analysis for CD19 combined cohort

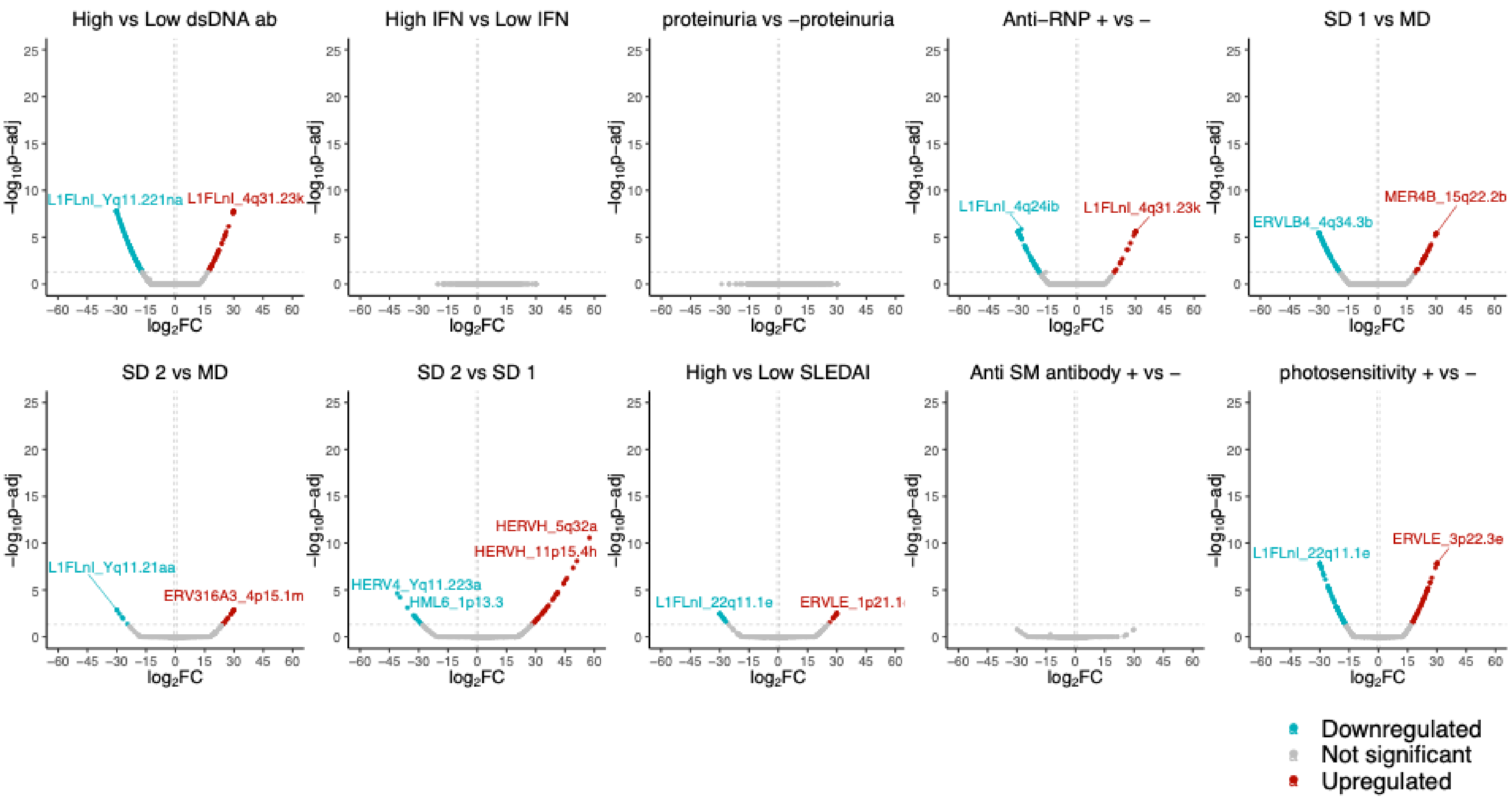


**Supplementary Figure 15.** SLE subphenotype differential TE analysis for CD19 european enriched cohort

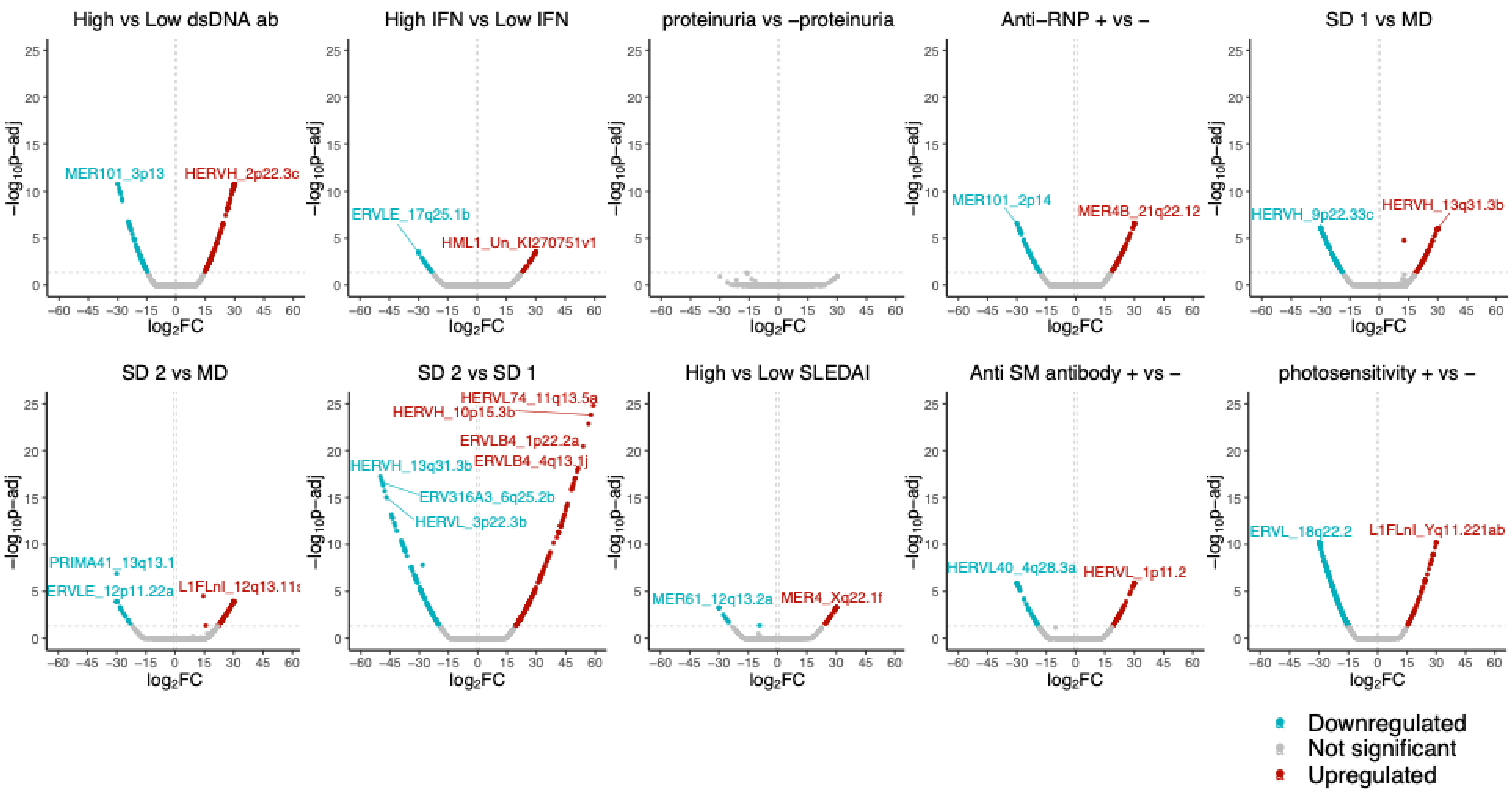


**Supplementary Figure 16.** SLE subphenotype differential TE analysis for CD19 asian enriched cohort

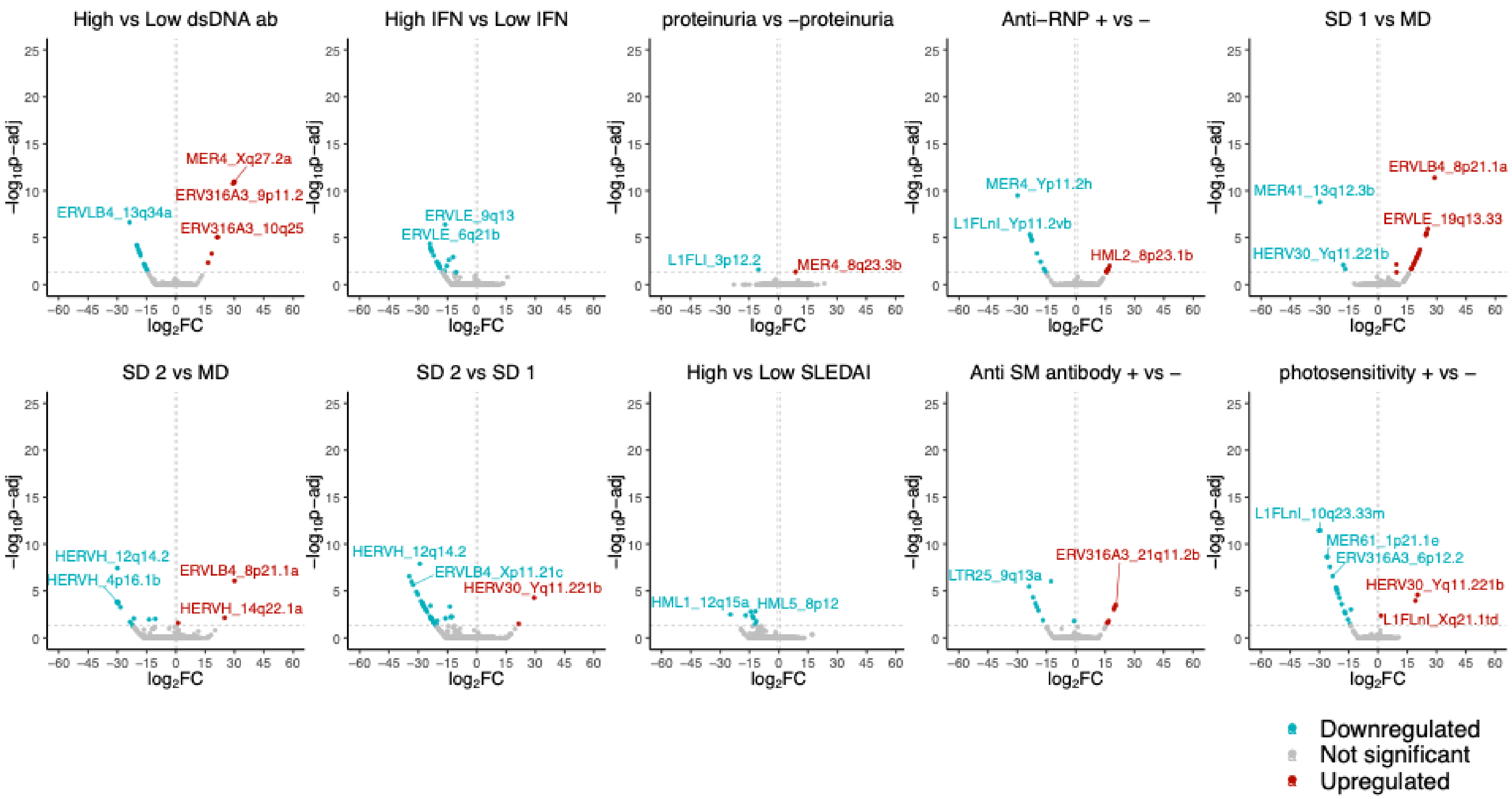


**Supplementary Figure 17.** SLE subphenotype differential TE analysis for NK combined cohort

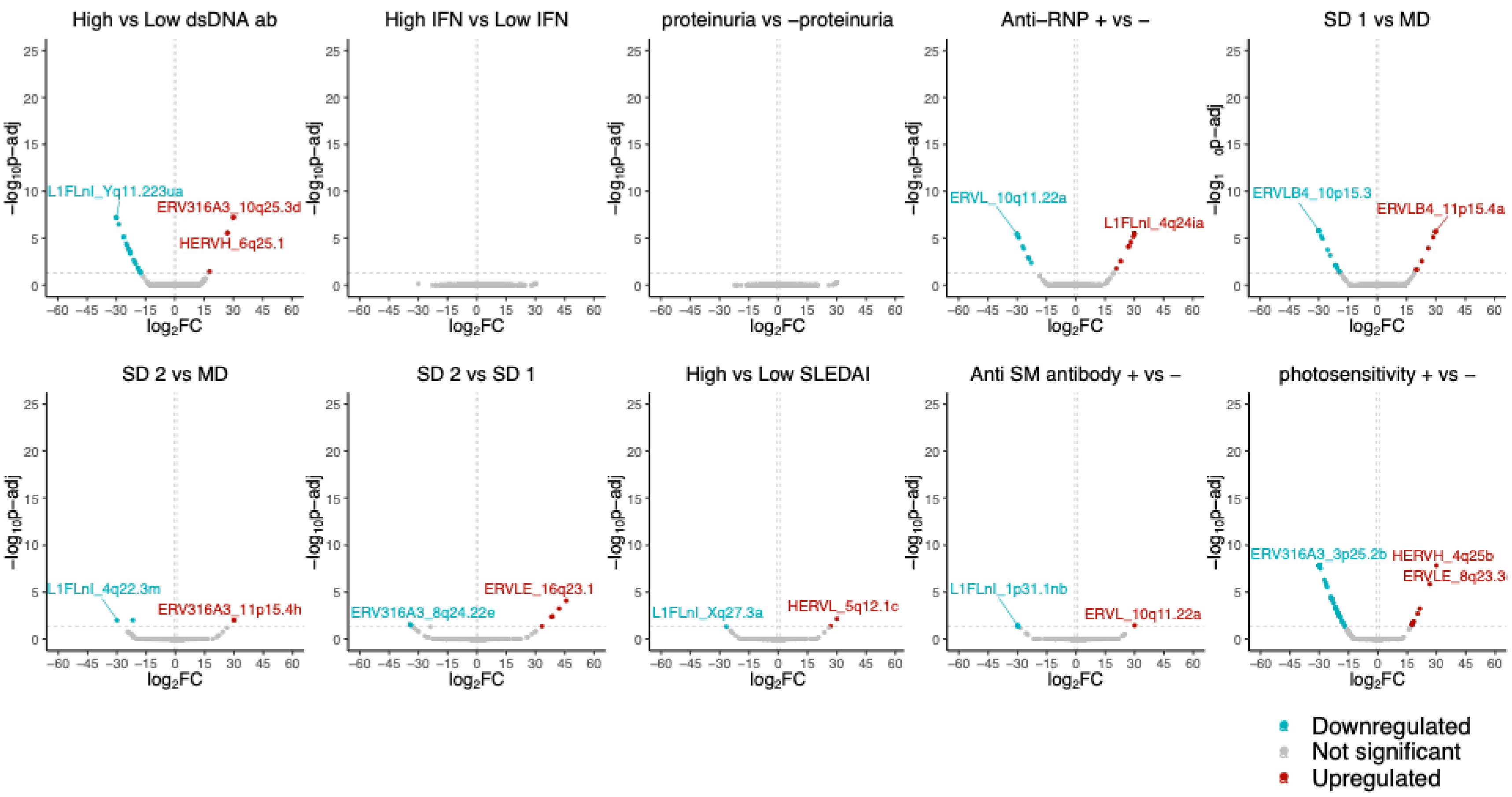


**Supplementary Figure 18.** SLE subphenotype differential TE analysis for NK european enriched cohort

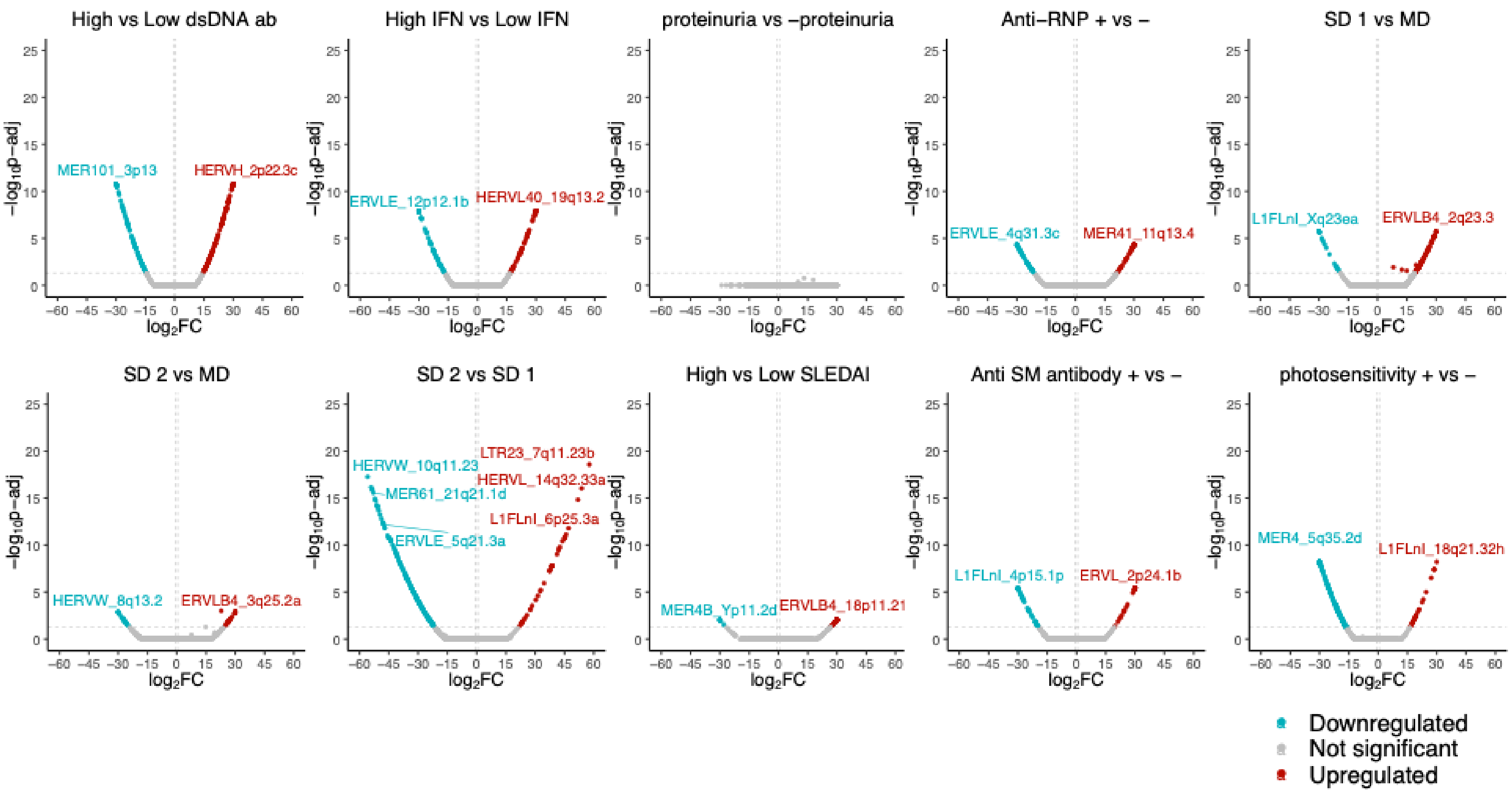


**Supplementary Figure 19.** SLE subphenotype differential TE analysis for NK asian enriched cohort
